# Unearthing the global impact of mining construction minerals on biodiversity

**DOI:** 10.1101/2022.03.23.485272

**Authors:** Aurora Torres, Sophus O.S.E. zu Ermgassen, Francisco Ferri-Yanez, Laetitia M. Navarro, Isabel M.D. Rosa, Fernanda Z. Teixeira, Constanze Wittkopp, Jianguo Liu

## Abstract

Construction minerals – sand, gravel, limestone – are the most extracted solid raw materials^1^ and account for most of the world’s anthropogenic mass, which as of 2020 outweighed all of Earth’s living biomass^2^. However, knowledge about the magnitude, geography, and profile of this widespread threat to biodiversity remains scarce and scattered^3–6^. Combining long-term data from the IUCN Red List and new species descriptions we provide the first systematic evaluation of species threatened by mining of construction minerals globally. We found 1,047 species in the Red List impacted by this type of mining, of which 58.5% are threatened with extinction and four species already went extinct. We also identified 234 new species descriptions in 20 biodiversity hotspots reporting impacts from mining. Temporal trends in the assessments highlight the increased saliency of this threat to biodiversity, whose full extent may well reach over 24,000 animal and plant species. While rock quarrying mostly threatens karst biodiversity and narrow-ranged species, sand and gravel extraction is a more prominent threat to freshwater and coastal systems. This study provides the first evidence base to support a global strategy to limit the biodiversity impacts of construction mineral extraction.

---

As of 2020, the human-made mass outweighs all of Earth’s living biomass^2^. Nearly 90% of such is made of construction minerals^7^, mainly aggregates (sand, gravel, and crushed rock) and limestone. These are the world’s most extracted solid materials by mass^1^, and their global market worth US$448.5 billion in 2020^8, 9^. Most of this mining has occurred since the 1980s due to rapid population, urban, and infrastructure growth^10^, and mining rates are predicted to double from 2017 to 2060^1^, dramatically intensifying pressures on biodiversity. Satisfying the demand for construction minerals without transgressing planetary boundaries represents an important and insufficiently recognized sustainability frontier^3, 11, 12^.

Mining of construction minerals (hereafter *construction mining*) occurs in every country, although developments grow faster in biodiversity-rich coastal zones and hotspots^12, 13^. Many source ecosystems (**Extended Data Fig. 1**) harbor endemic species and highly-diverse communities that are crucial for ecosystem functioning and services supply, including food and clean water provision, and land stability^14, 15^. Mining poses serious, often irreversible and far-reaching impacts, to those ecosystems, for example through erosion, shrinking deltas, salinization, pollution, and traffic disturbances^6, 16–20^. Despite this, the global biodiversity burden associated with construction mining remains understudied^4–6^ and the indirect impacts of infrastructure on biodiversity through materials sourcing are rarely assessed^21^. A lack of global mining datasets of construction minerals (barely represented in SNL Metals and Mining or S&P Global Market Intelligence databases), and reporting challenges of a mostly domestic and often largely informal extractive sector^11, 22^, make quantifying this threat a daunting challenge.

Here, we present the first global evaluation of the impact of construction mining on species endangerment. We used two complementary information sources: the IUCN Red List of Threatened Species (hereafter *Red List*) and new species descriptions. We conducted a systematic search through the 75,530 assessments with threat description by 2020^23^ for the former and peer-reviewed literature for the latter, resulting in 1,082 Red List assessments and 141 studies describing new species reporting threats from construction mining (hereafter simply *impacted by construction mining*). We used these data to determine (1) how many known species are impacted by construction mining and what are their temporal trends; (2) where are higher levels of threat reported; and (3) which types of species are most likely to be impacted.

## Magnitude of threats

Overall, construction mining has contributed to four known species extinctions and impacted at least 1,047 species and 19 intraspecific taxa assessed by the Red List by 2020, of which 58.5% are threatened with extinction (classified as: Critically Endangered; Endangered; or Vulnerable) (**Figs. 1A-B**, see **Extended Data Supplementary Results**). Additional 234 new species descriptions reporting threats from construction mining were identified, of which only 10.7% are in the Red List, mostly as threatened (80%) and Data Deficient (12%). Half of all identified Red List taxa are impacted by sand and gravel extraction (51.1%) and the other half by rock quarrying (48.9%), predominantly of limestone (81.4%). Two thirds of the new species (67.1%) are impacted by rock quarrying mostly of limestone (96%).

**Figure 1.**
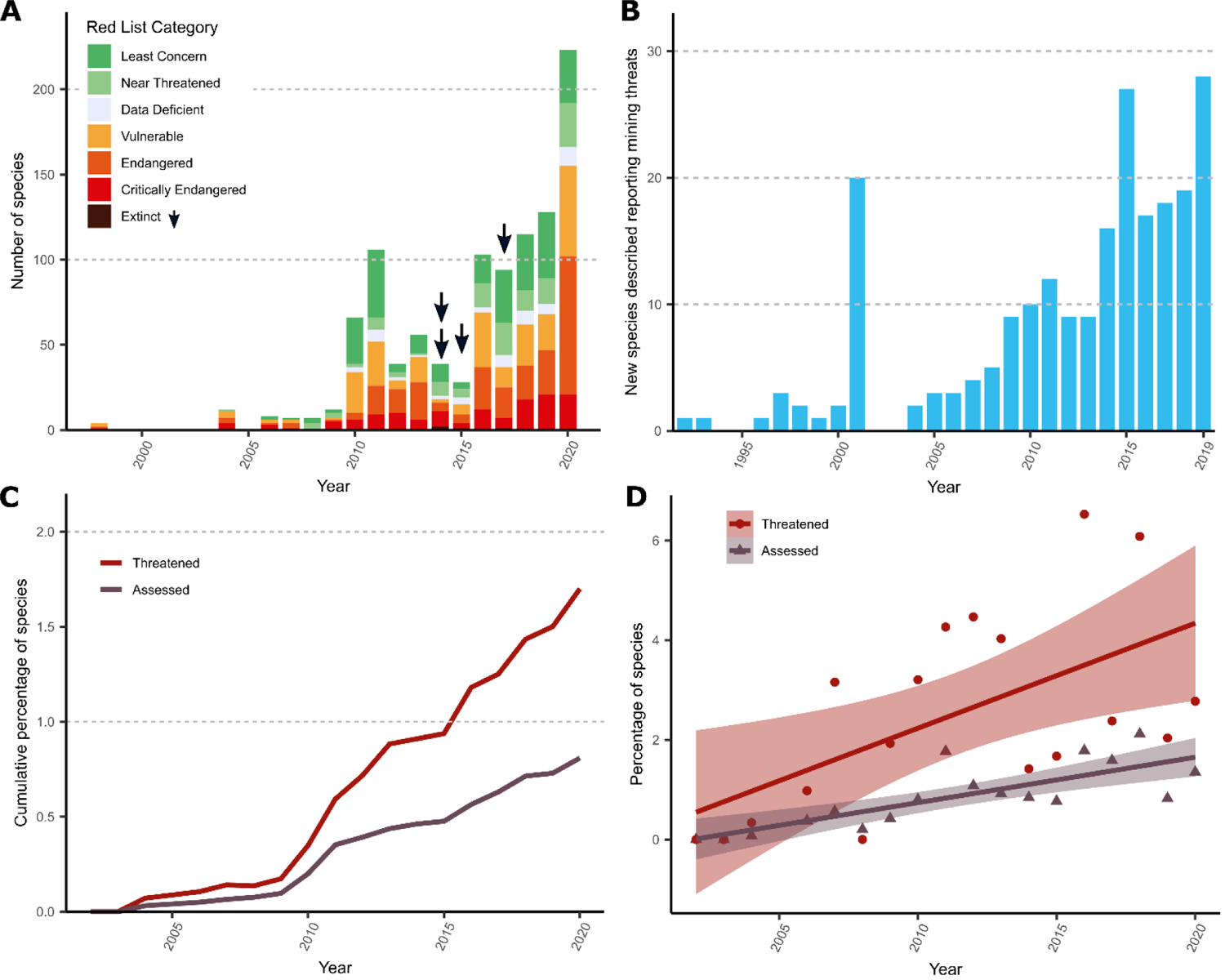
Temporal trend of construction mining threat reports indicating that this pressure is becoming a salient threat. (**A**) Number of species added to the Red List per year impacted by construction mining, classified by their conservation status. Arrows indicate extinction events. (**B**) Number of new species descriptions per year reporting impacts by construction mining. (**C**) Cumulative percentage of species assessed (all Red List categories) and those threatened with extinction (Critically Endangered, Endangered or Vulnerable) of the Red List impacted by construction mining over time. (**D**) Increase in the percentage of species impacted by construction mining of the total species assessed in the Red List per year (*P* < 0.05).

Red List assessments and new species descriptions reporting impacts from construction mining have grown over the last decade (**Fig. 1A-B**). An increase in absolute and relative level of threat, reaching 1.71% and 0.83% of the total number of Red List threatened and assessed species in 2020, respectively (**Fig. 1C-D**), and the growing number of species descriptions reporting impacts (*P* < 0.001) despite a stable trend for new descriptions^24, 25^ suggest that construction mining is an increasingly salient threat to biodiversity.

## The geography of threats

Red List species impacted by construction mining are mainly distributed in a ‘belt’ from Southeast Asia towards the Middle East, Northern Africa and Western Europe, and particular locations in the Congo River, Guinea, and Brazil’s Atlantic Forest and Cerrado (**Fig. 2A**), including biodiversity hotspots such as Wallacea, Sundaland, Indo-Burma, Himalaya, Western Ghats, and the Mediterranean basin as identified by Myers *et al.*^26^ and Conservation International^27^. The type localities of 72.6% of the new species impacted by construction mining were found within 20 biodiversity hotspots, predominantly in Indo-Burma (**Fig. 2B, Extended Data Fig. 4**), greatly overlapping those and other hotspots poorly captured by the Red List assessments such as California, Succulent Karoo, Cape floristic province, and Coastal forests of Eastern Africa. While 93% of descriptions up to 2001 were reported in the US, New Zealand, and Australia (**Extended Data Fig. 5**), after 2010 over 70% came from India, Brazil, Malaysia, China, and Vietnam, coincident with rapid urban and infrastructure expansion^12^. Overall, these patterns are very different from those of mines of metals and precious minerals^19, 28^, except in a few coinciding areas such as the Iron Quadrangle in south-eastern Brazil.

**Figure 2.**
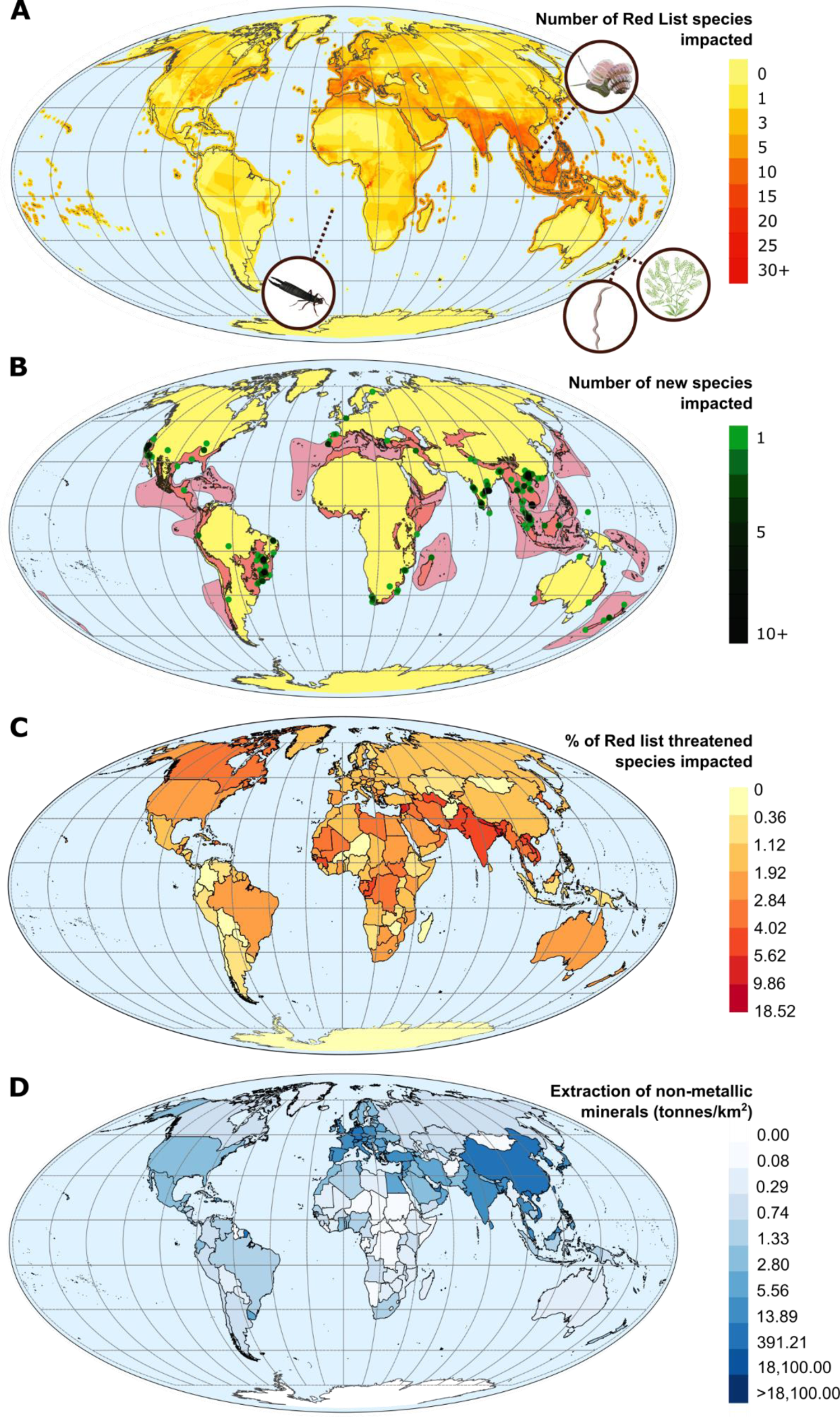
Global spatial patterns of species impacted by construction mining. (**A**) Distribution ranges of the identified Red List species with spatial data (*n =* 824), including the extinction location of 4 species: Saint Helena giant earwig (*Labidura herculeana*); Schmarda’s worm (*Tokea orthostichon*); a peppergrass (*Lepidium obtusatum*); and a land snail (*Plectostoma sciaphilum*) (see **Extended Data Fig. 2** for the distribution of only species threatened with extinction). Every individual species is given equal weight, regardless of its threat level. The actual impact might vary across their range. As virtually no construction mining occurs in the open oceans, the sea surface beyond the continental shelf and at a distance >100 km from the coastline was excluded for clarity. (**B**) Type localities of new species impacted when described (*n =* 234) across global biodiversity hotspots^27^. (**C**) Percentage of Red List threatened species impacted by construction mining over the total number of Red List threatened species per country (**Extended Data Fig. 3** for assessed species). (**D**) Domestic extraction of non-metallic minerals (construction dominant) by area from UN and IRP Global Material Flows Database^30^.

The ranges of the identified Red List species concur partly with the amount of extraction of construction minerals per country (**Fig. 2A, C-D**). We hypothesized that countries that have undergone higher levels of economic growth, construction minerals extraction, and with weaker governance would have a higher proportion of species impacted by construction mining. Beta-regression models show that the world region was the most significant predictor of both the proportion of assessed and threatened species impacted by mining per country, followed by the Corruption Perception Index (CPI) (**Extended Data Tables 1-2**), indicating strong geographical patterns and that countries with stronger governance have a lower proportion of impacted species, as suggested by previous publications^29^. A higher proportion of assessed species impacted by construction mining is found in Middle East and North Africa, followed by Western Europe, Eastern Europe-Central Asia, and Asia-Pacific (**Extended Data Figs 7-8**). The proportion of Red

List threatened species was also highest in the Middle East and North Africa, followed by the Asia-Pacific region. Indeed, the countries with the largest percentage of threatened species impacted by construction mining are Lebanon (18.5%), Trinidad and Tobago (9.9%), Bangladesh (7.3%), Syria (7.3%), and Nepal (7.0%) (**Fig. 2C**). Proxies for the intensity of construction mining such as the volume of mineral extraction, or GDP per capita did not significantly correlate with the proportion of assessed or threatened species.

## Taxonomy, habitats, and mining patterns

Our analyses revealed distinct associations between mining threats, other threats, and habitat types, providing insights into the characteristics of the species impacted by construction mining. Key threat associations with construction mining-reported threats were the presence of urban, transport, and energy infrastructure, pollution, human intrusions, and natural systems modifications (**Fig. 3A**). Overall, species from freshwater (e.g., river, deltas), coastal habitats (e.g., dunes, seagrasses), and cave and rocky areas are more likely to be described as impacted by construction mining (**Fig. 3B**), coinciding with some of the most threatened ecosystems at present^16, 31^. Dicots, fishes, gastropods, and reptiles featured more prominently (30.3%, 13.2%, 10.3%, and 8.2%, respectively) across taxonomic groups (**Fig. 4**). Adding the data on species descriptions broadened the taxonomic scope of impacts towards arthropods, vascular plants, bryozoans, and ascomycotan (**Extended Data Fig. 9**). An interactive diagram with the full taxonomic details of the species included in the **Supplementary Information**.

**Figure 3.**
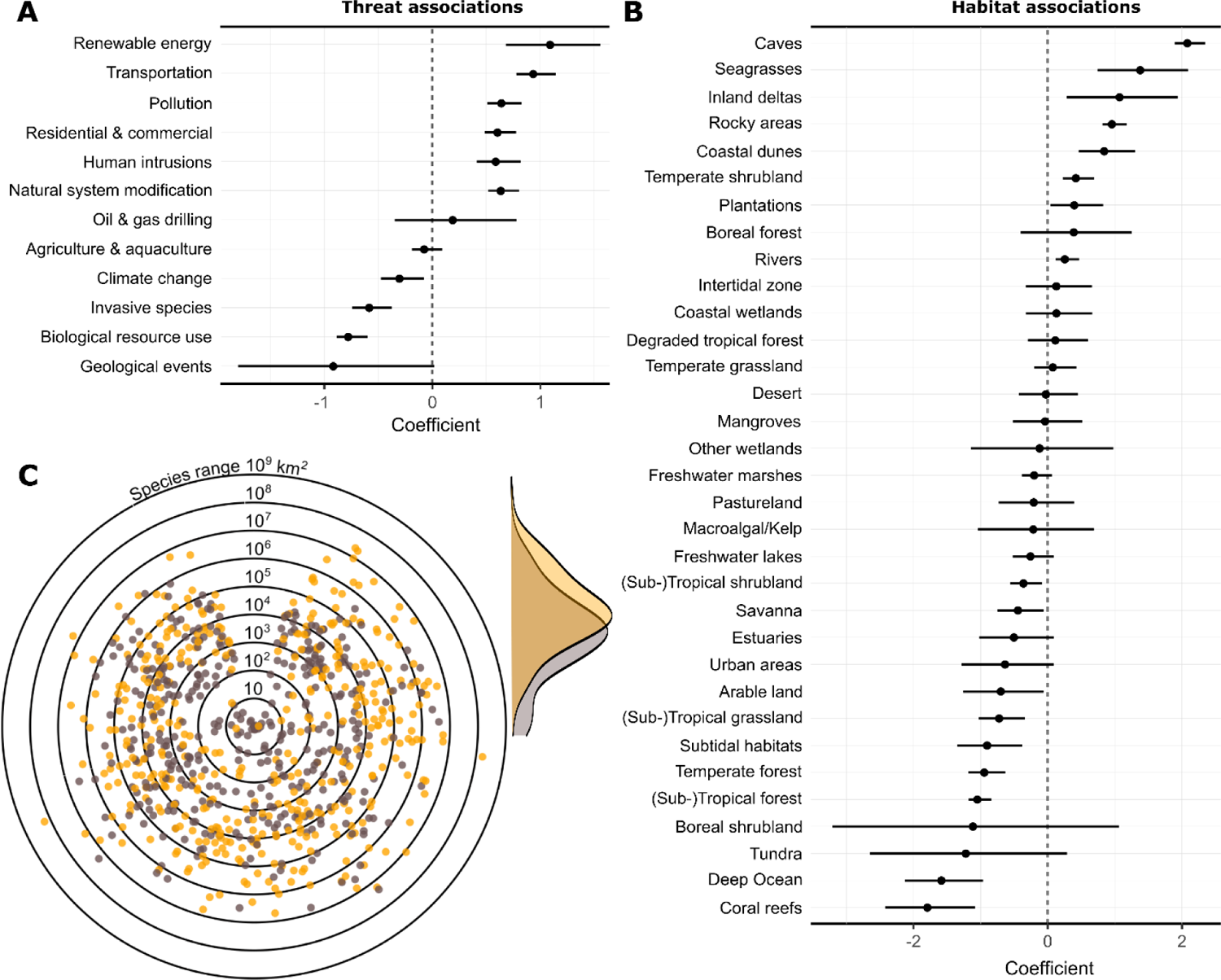
Characterization of species impacted by construction mining in relation to habitat associations, threat associations, and their geographic range. (A) Threat and (B) habitat associations of species impacted by construction mining when compared to all Red List assessed species. Coefficient estimates and their 95% confidence interval for the binomial logistic regression between being impacted by construction mining and habitat types (**Supplementary Table 1**) and other threats. (C) Documented species and mining types in relation to geographic range (sand and gravel extraction in yellow and stone quarries in grey). Each dot indicates a species, located randomly along the perimeter of a circle with radius equal to the log10 of the species’ range size in km^2^. The density plots of range observations from 10 to 10^9^ km^2^ are shown on the side.

**Figure 4.**
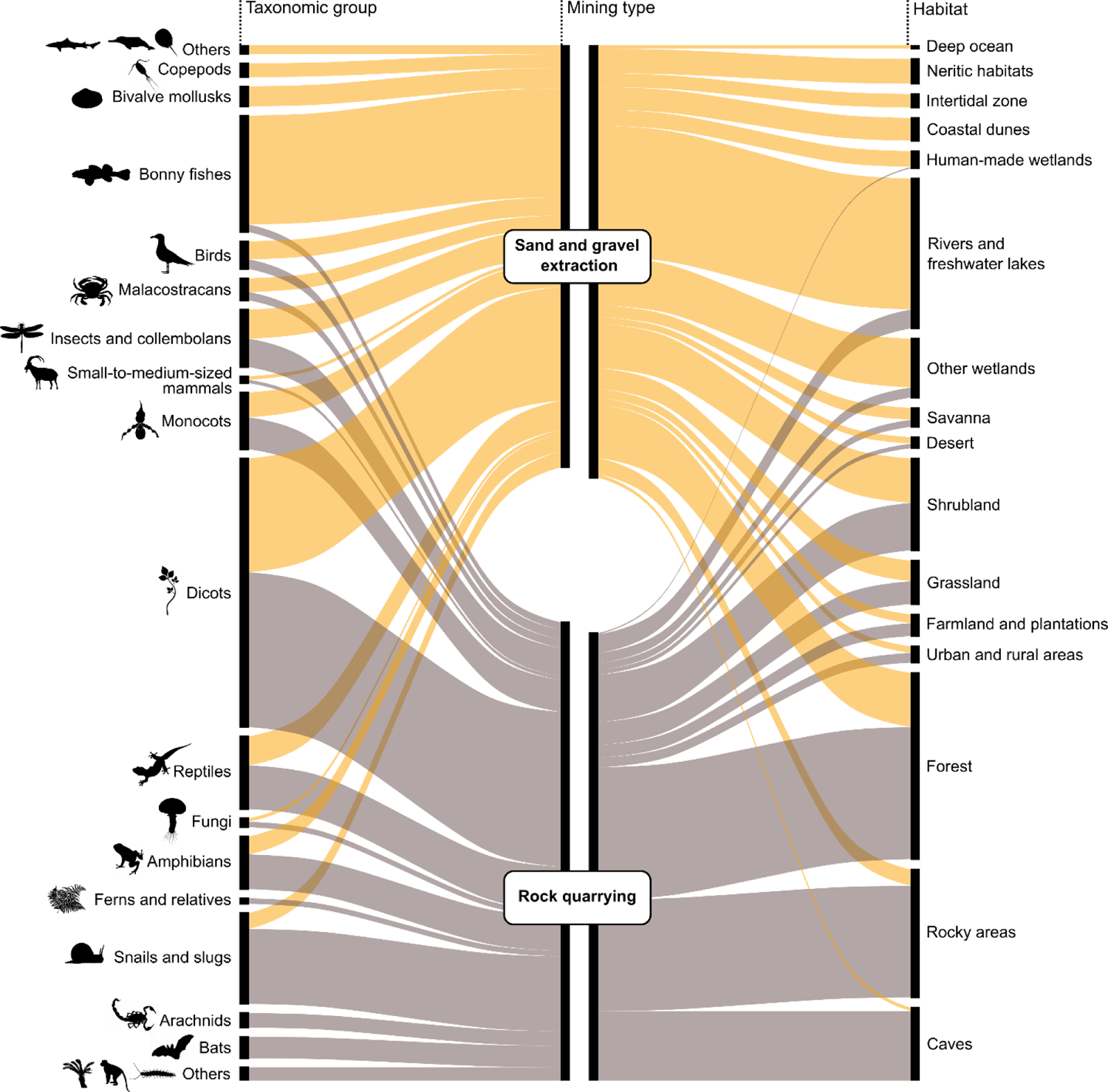
Taxonomic group and habitat in relation to mining type for species impacted by construction mining in the Red List and species descriptions (*n* = 1,275). The width of each box is proportional to the number of species and intraspecific taxa included per group, mining type, and habitat type (**Supplementary Table 1**), while colors denote mining type. Groups are sorted by the proportion of records under each mining type. ‘*Others’* taxonomic groups (with < 10 observations) impacted by sand and gravel extraction include: sirenians; porpoises, and dolphins; sharks and rays; bryozoans; horseshoe crabs; conifers. ‘*Others’* taxonomic groups impacted by rock quarrying include: primates; millipedes; mosses; cycads; corals. Groups with 1 observation are not shown (Annelids: one species threatened by sand extraction, and one species by rock quarrying). Icon credit—http://phylopic.org. Icon Attributions licensed under CC BY-SA 3.0: fern by Olegivvit, dragonfly and scorpion by Gareth Monger, dolphin by Chris huh, crab by Hans Hillewaert (vectorized by T. Michael Keesey), millipede by Ralf Janssen, Nikola-Michael Prpic & Wim G. M. Damen (vectorized by T. Michael Keesey).

There was substantial separation among impacted taxonomic groups, habitats, and their range sizes by mining types (**Figs. 3C** and **4**). Reported threats from sand and gravel extraction dominated on species of rivers, wetlands, and coastal habitats, with affected taxa ranging from freshwater and marine megafauna (e.g., river dolphins, crocodiles) to the meiofauna of hyporheic and groundwater habitats. Sea dredging also posed a significant threat to species inhabiting seagrass meadows. In contrast, reported threats from rock quarrying were more common among species of dicots, gastropods, bats, amphibians, monkeys, arachnid, fern and cycads that occur in forests, rocky habitats, and caves, many of which are associated with karsts.

Small geographic ranges dominated across impacted species, with 50% of species having ranges < 10^4^ km^2^. Species impacted by rock quarrying have significantly lower range sizes (median [IQR] range size: 4,184 km^2^ [419-25,387]) than species impacted by sand and gravel extraction (median [IQR] range size: 22,963 km^2^ [4,015-174,560]) (*F*-statistic = 78.58, *P* < 0.001), potentially owing to easier propagation of mining impacts in river and coastal ecosystems^32^, a more partial reliance on sand-gravel habitats for certain stages of lifecycle (e.g., as nesting habitats, nursery and feeding grounds, or migratory stepping stones)^33, 34^, and limited dispersal abilities of karsts and cave species, including relict or highly specialized species^16^.

Growing environmental concerns about river mining are contributing to increasing aggregates production in quarries^11, 35^. Our results, however, show that such transition does not necessarily reduce species extinction risks, particularly when quarrying occurs in poorly documented areas and/or with high levels of endemism^36^ where entire communities can become extinct^4,16, 37^. Greater appreciation of the diversity of mining contexts and mineral-supply networks is needed to quantify and characterize the impacts of extractive activities and to integrate them in the supply-chains of urban and infrastructure development projects.

## Global estimate of threatened species

Our work provides a baseline of known species threatened by construction mining that is likely to be just the tip of the iceberg. Only 6% of the world’s described species have been assessed by the IUCN, with groups such as invertebrates, fishes, plants and fungi – which account for 78% of our database – being particularly underassessed^38^. Moreover, many species remain undiscovered^39, 40^, and many will have gone extinct even before description^41^. We used simple extrapolation methods used by the Intergovernmental Science-Policy Platform on Biodiversity and Ecosystem Services^42, 43^ to estimate how many of the planet’s species are threatened (see **Methods**). On average, 0.50% of the Red List threatened animal and plant species (excluding insects) and 0.21% insect species are impacted by construction mining (**Supplementary Table 4**). Extrapolating by the number of estimated extant species results in an estimate of 24,246 species [24,234-27,226]. This first estimate should be regarded as a conservative indication of the order of magnitude of the total number of threatened animal and plant species.

Indeed, our results suggest the likely under-recording of construction mining as a threat. First, the proportion of species impacted by construction mining in the Red List is growing rapidly every year (**Fig. 1C-D**). Second, the number of Red List species impacted by construction mining is not commensurate with the scale of the activity, with only a tenth of the assessed species impacted by *mining and quarrying* being explicitly impacted by construction mining. Third, the lack of relationship between the volume of extraction of construction minerals and the proportion of Red List species impacted by construction mining points at extensive and uneven underreporting, further evidenced by local threat clusters. For example, China produces 44.3% of the world’s construction aggregates^45^ and consumed in three years more cement than the US did in the entire 20^th^ century^46^. Yet the number of threatened species impacted by mining in Lebanon is almost twice as high, despite having 0.1% of China’s area (22 to 40 species in China and Lebanon, respectively). Finally, construction mining might be masked by co-occurring or linked threats.

Most extraction occurs close to building sites^11^ (**Fig. 3**), yet *residential and commercial development* is classified as a threat on the Red List almost 20 times more frequently than construction mining. It is important to acknowledge that active, abandoned or restored pits and quarries may contribute to the conservation of rare and threatened species and habitats (e.g., sand martin *Riparia riparia,* flamingo moss *Tortula cernua*)^47–49^, although opportunities are context- and site-dependent^48, 50^ and there is a severe evidence gap on how impacts can be offset to achieve no net loss^51, 52^. Considering all these points, and the time-lags in recognizing threats and extinction events affecting the Red List^53, 54^, the real number of species impacted could be much higher.

## Conservation and restoration implications

Rapid action to reduce mining pressures and restoring habitats and populations will contribute to ameliorate this threat. Our assessment flags threats for neglected groups of above and below ground organisms^36, 37, 55^. Mining operations must improve implementation of the mitigation hierarchy setting, avoid non-offsettable impacts (Pilgrim *et al.* 2013; Simmonds *et al.* 2021), and improve the evidence on effective offsetting unavoidable impacts. When expanding into areas where species are threatened or biodiversity remains poorly documented, extractive industries should follow a risk-based approach^58^. An undue focus on the Red List has led the impact assessment process to safeguarding the most threatened species, whereas vulnerable, unassessed or poorly assessed species easily become forgotten^59, 60^. We call for setting good practice in impact assessment so that if a species is potentially new to science or globally threatened, has highly restricted range, and knowledge of its distribution, ecology, and of restoration needs is lacking, the precautionary principle should apply and impacts on it should be avoided (Treweek pers. comm.). If all actors decide avoidance is unfeasible, it should not be translocated, moved or destroyed until its requirements are researched and effective techniques are available.

## Conclusions

Our results quantitatively demonstrate that during the last decade construction mining has emerged as a global environmental challenge^5, 61^. This assessment shows that construction mining represents a growing threat that affects biodiversity of land, freshwater and coastal systems alike, understudied taxonomic groups and range-restricted species. The magnitude of this threat will continue to grow along with urban and infrastructure development^1, 32^, with an overwhelming risk of prioritizing developments that foster economic expansion over conservation objectives, compromising our ability to halt biodiversity loss and achieve nature-focused Sustainable Development Goals^62, 63^. Mobilizing resources and efforts to mainstreaming biodiversity in the mining and construction sectors should become urgent priorities and key actions to implement the upcoming post-2020 Global Biodiversity Framework.

## Supporting information

Supplementary Tables 1-4 and Additional Results

## Methods

### Data search

To build a first database of species impacted by construction mining we used two complementary data sources: the IUCN Red List of Threatened Species and new species descriptions. The detailed Red List assessments document the threats affecting species and are the most comprehensive and widely used information source on the conservation status of species globally^64^. Given the Red List taxonomic incompleteness and its difficulties capturing emerging threats and extinction events^54, 65, 66^, we also examined how construction mining is reported in the description of new species and how their taxonomic and geographic coverage differs from those of Red List species. Data processing and statistical analyses were carried out in the R statistical software v. 4.0.3^67^.

We searched through the Red List of Threatened Species version 2020-3^23^ to identify and characterize relevant assessments reporting threats from construction mining by 2020. Although a category of threats exists for *mining and quarrying* this subset includes a wide variety of minerals (e.g., gold, diamonds). Our main focus was on the mining of the world’s most extracted construction minerals^1^: aggregates and limestone. We searched through the 75,530 individual Red List assessments with threat description available, including global and regional assessments (Arabian Sea, Caribbean, Central Africa, Eastern Africa, Europe, Global, Gulf of Mexico, Mediterranean, Northeastern Africa, Northern Africa, Pan-Africa, Persian Gulf, S. Africa FW, Western Africa), using a text-mining script in R to identify potentially relevant assessments through a search of keywords (both with lowercase and capital letters): sand, gravel, limestone, cement, stones, aggregate, concrete, quarr, dredging. We identified 2,027 assessments for screening and data extraction.

Screening and coding of Red List assessments was performed by four reviewers (SOSEzE, CW, FF, AT). A subset of 10% of all assessments was screened by two reviewers (Kappa test: *k* > 0.75), with all disagreements discussed in detail. Refinements of the inclusion criteria were made in liaison with the entire review team where necessary.

The following inclusion criteria were used to assess the relevance of assessments identified through searching:

- Relevant subjects: Species, subspecies, varieties or subpopulations. No geographical or taxonomic restrictions were applied.
- Relevant types of intervention: Described threat from mining, quarrying, and/or dredging of construction minerals – namely, construction aggregates (sand, grave and crushed rock), silica sand for glass production, limestone, other typical rocks used in construction either for aggregates production or as dimension stones (e.g., sandstone, slate, marble, granite, basalt, gypsum). We included every kind of mining operation, which can range from industrial mining to artisanal mining activities and those of a legal, illegal or informal nature. The link with construction is assumed from the type of mineral mentioned (see list above), explicitly mentioned in the assessment, or inferred from the description of the threat (e.g., mention of quarrying plus urban or road construction). Finally, assessments must refer to the mining stage and not the restoration or abandonment stage.

When the eligibility of assessments was unclear, the authors of assessments were contacted to provide additional details. For all excluded assessments, reasons for exclusion were provided in the form of one or more a priori exclusion criteria as follows:

- “Exclude, not mining”. The species is not relevant for the assessment as mining is not mentioned in the threat description.
- “Exclude, material not mentioned”. Mining or quarrying is mentioned as a threat but it does not specify the mineral/rock that is quarried nor the intended use.
- “Exclude, not relevant mining”. Mining of precious minerals (diamonds), metals (e.g., iron) or fuel (e.g., oil, tar or bituminous sands) were not deemed relevant for this study.
- “Exclude, not relevant mining phase”. Assessment exclusively highlighting the impacts associated with restoration or recolonization of mine sites were excluded.

The total number of relevant Red List assessments post-screening was 1,082. For these assessments, we downloaded all the taxonomic, geographic, habitat, and threat information from the Red List database, and we coded mining-related variables based on the information provided in threat assessments: mineral mined (e.g., aggregates from unconsolidated deposits, rocks); mining method (e.g., dredging and excavation; blasting and quarrying); mining type (e.g., sand and gravel extraction; rock blasting and quarrying).

To identify new species descriptions reporting threats from construction mining we implemented searches in Web of Science (13 bibliographic databases), Scopus, and Google Scholar in November 2020. All bibliographic databases were accessed using Michigan State University’s institutional subscriptions. For the searches in Google Scholar, we used three simplified search strings (on sand, gravel, and limestone) to search for additional species descriptions. The cut-off of relevant hits for each search string was determined through a scoping exercise (see **Supplementary Information File 1**). The search results were exported for screening in an Excel spreadsheet and duplicates were removed. Authors of publications unobtainable through library licenses were contacted to request access to the full article. The reference sections of reviewed articles and any relevant reviews found during searching were manually searched to identify additional articles. Eligibility screening and coding of papers was conducted at full-text and performed by a single reviewer (AT), following the inclusion/exclusion criteria mentioned above. A total of 1,581 papers were identified and reviewed in full. Of those, 141 papers included relevant information and were coded. Of the excluded papers, 20 papers were not accessible (no available pdf), 50 papers did not mention the specific mineral being exploited, 68 papers reported threats from other mining types (e.g., diamonds, gold), and 1,302 papers were irrelevant. The following data from new species descriptions were coded (Meta-data extraction and coding schema are available in the **Supplementary Information File 1**): publication characteristics, taxonomic information, proposed red list category, red list criteria, mineral mined, mining type, mining method, legality of mining, number of threats, intensity of mining threat, timing, rationale of threat, habitat type, and geographic location.

### Spatial analyses

To assess geographical patterns in mining threats, we mapped the distribution of species impacted by mining of construction minerals. We compiled the range maps for identified species with available spatial information from the IUCN^23^, and for birds from Birdlife International^68^ (*n* = 823 species in total). Processing and mapping were done in QGIS desktop version 2.18.17^69^, Python, GDAL library and R software. The retrieved data consisted of polygons in shapefile format and point data in CSV format. The geographic range size was calculated after re-projecting to a global equal area projection (Molleweide, datum WGS84). For point data, we used a minimum convex polygon to calculate the range size or extent of occurrence of each species following IUCN Mapping Standards version 1.19 (https://www.iucnredlist.org/resources/mappingstandards). To avoid having species with a distribution area of 0 km^2^ and to prevent a significant number of the locations to falling on the edge of the distributional range of the species, point data was first converted into polygons by applying a 10 km buffer to each point. A convex hull was then applied around those 10 km buffers. As virtually no mining of construction minerals occurs in the open oceans, all the sea surface beyond the continental shelf or at a distance > 100 km from the coastline was excluded from the maps for visualization purposes (**Fig. 2A**). For the representation of the range size data (**Fig. 3C**), the range size for each species was located randomly along the perimeter of a circle with radius equal to the log10 of the species’ range size in km^2^, differentiating by mining types (sand and gravel: *n =* 418; rocks: *n* = 405). As only one species was impacted by both sand mining and limestone quarrying (*Margaritifera margaritifera*), we re-coded its mining type to sand and gravel extraction for visual representation, as sand mining is a more direct pressure to freshwater mussels. Using country-level data from the Red List, we calculated the proportion of species (including intraspecific taxa) impacted by construction mining for each country from the total of Red List species assessed and threatened for each country, excluding all uncertain distributions, introduced species and vagrant records.

The type localities of newly described species were extracted from their associated publication. When coordinates – latitude and longitude – were not directly provided we retrieved them by checking GBIF (https://www.gbif.org/), georeferencing maps of the original publication or, when the latter was not provided, by using locality annotations and Google Earth Pro to identify the approximate location.

### Modelling country-level threat patterns

To explore whether any country-level characteristics explained differences in construction mining-related threats to species, we ran exploratory regression models. Our response variables were the proportion of species assessed and threatened with extinction reported as impacted by construction mining of the total number of assessed and threatened species in the Red List by country. We fitted beta regression models with a logit link function using the *betareg* function of the package ‘betareg’^70^. The significant threshold was *P* < 0.05. Explanatory variables hypothesized to affect the outcome were the same in both models: 1) world region from the World Bank data^71^; 2) GDP per capita in the most recent year for which there was data available (all between 2017-2020) calculated using GDP and population size data from the World Bank data^71^; 3) average domestic extraction of construction minerals 2010-2017 per hectare, calculated from the Global Material Flows Database of the UN International Resource Panel^30^ http://www.resourcepanel.org/global-material-flows-database using “Non-metallic minerals - construction dominant” flows; and 4) Transparency International’s Corruption Perceptions Index (CPI) for 2020, which takes values from 0 (highly corrupt) to 100 (very clean) and was downloaded from https://www.transparency.org/en/cpi/2020/index/nzl. The CPI is a metric of the perceived public sector corruption within a country based on data collated from surveys of experts and business people and is a widely used proxy for strength of national governance^72^. The above data was available for 169 countries. We tested alternative variables of domestic extraction of construction minerals from the Global Material Flows Database: total extraction in 2017 and average extraction 2010-2015 per capita (**Extended Data Fig. 6**), but they did not alter model performance. We acknowledge that other factors that might be linked with the proportion of assessed or threatened species such as habitat amount or the degree of red-listing effort per country were not included in the model. However, the broad diversity of habitats affected ranging from intensive human land-uses to natural habitats makes difficult to control by this factor. Both models were evaluated by analyzing the residuals and showed reasonable goodness of fit (**Extended Data Fig. 8A-B**). The model with the proportion of assessed species explained a higher amount of the variance (Log-likelihood = 817.4, *P* < 0.001, Pseudo *R*^2^ = 0.60) than the model with the proportion of threatened species (Log-likelihood = 500, *P* < 0.001, Pseudo *R*^2^ = 0.14), but both were similar in significance, direction and magnitude. Tukey’s post-hoc tests were employed for pairwise comparisons among world regions using the ‘emmeans’^73^ package (**Extended Data Fig. 8C-D**).

### Habitat associations

To investigate the relationship between species habitat and whether the species is impacted or not by mining of construction minerals we carried out a generalized linear model using the *glm* function, family binomial, and a logit link function. We used all species assessed in the Red List database with reported threats, excluding Data Deficient species (*n =* 44,463, of which 1,003 are reported as impacted by construction mining). When multiple assessments were available for a given species (e.g., European assessment and Global assessment in *Testudo graeca*), but only one of the assessments reported threats, the other assessments were excluded from the analysis. This situation affected 7 species: *Testudo graeca, Potamogeton schweinfurthii, Cyclosorus interruptus, Apterichtus anguiformis, Apterichtus caecus, Trachyrhamphus bicoarctatus*, and *Alaudala rufescens*. The response variable represented whether a species was impacted (or not) by construction mining and the predictor variables were occurrence (or not) in each habitat type. We started by testing the first level habitat types (16 types) from the IUCN Habitats Classification Scheme version 3.1 (https://www.iucnredlist.org/resources/habitat-classification-scheme). However, these classes were too broad for interpretation. We then tested second level habitat types (109 habitat types), but the model showed a lack of fit. Then, we decided to combine these 109 habitat types into 33 categories (**Supplementary Table 1**). We assessed model fit using the Hosmer & Lemeshow goodness-of-fit test^74^ with the *hoslem.test* function from the ‘ResourceSelection’^75^ package. The Hosmer-Lemeshow test results (*χ*^2^= 9.785, *df* = 8, *P* = 0.280) indicated an overall goodness of fit of the model.

### Threat associations

To assess associations between construction mining-impacted species and other threat types on the Red List, we carried out a generalized linear model using the *glm* function, family binomial, and a logit link function. We used all the downloaded global assessments in the Red List database with available threat information and all assessments of species impacted by construction mining, excluding Data Deficient species. For those impacted species with multiple assessments, we kept the assessment that identified the construction mining threat and if multiple assessments remained we excluded non-global assessments. The response variable represented whether a species was impacted (or not) by construction mining and the predictor variables were being impacted (or not) by other 13 broad threats. For the threat types, we used the first level hierarchy of the IUCN Threats Classification Scheme version 3.2 (https://www.iucnredlist.org/resources/threat-classification-scheme). As *mining and quarrying* is found in the class *Energy production & mining* we used the level-2 hierarchy of this threat, to separate mining from the threats *Oil and gas drilling* and *Renewable energy*. We excluded species reported as impacted by *Mining and quarrying* (threat classes 3 *Energy production and mining* and 3.2 *Mining and quarrying*) but not specifically by construction mining to avoid considering as non-impacted species that might be impacted but whose threat description did not specify the mined minerals. We also excluded the class *Other threats* for being unspecific. In total, we examined associations for 53,519 species, of which 914 were impacted by construction mining. We assessed model fit using the Hosmer & Lemeshow goodness-of-fit test^74^ with the *hoslem.test* function from the ‘ResourceSelection’^75^ package. The Hosmer-Lemeshow test results (*χ*^2^= 21.149, *df* = 8, *P* = 0.007) indicate a poor goodness of fit of the model.

### Alluvial diagrams of taxa and habitat

Alluvial diagrams relating taxonomic group and habitat with mining type for species impacted by construction mining (*n =* 1,275 of which 1,066 are Red List species and intraspecific taxa and 209 are new species not in the Red List) were built using the R packages ‘ggplot2’^76^ and ‘alluvial’^77^ and the Inkscape software^78^. Taxonomic groups were organized into 19 classes: fishes; snails and slugs; bivalve mollusks; bats; malacostracans; small and medium-sized mammals; reptiles; insects and collembolans; amphibians; annelids; dicots; monocots; ferns and relatives; arachnids; birds; fungi; copepods; others (sand); and others (rock). The ‘*others*’ classes combined taxonomic groups with less than 10 observations. Those threatened by sand and gravel mining included: sirenians, porpoises, and dolphins (6), sharks and rays (3), bryozoans (2), horseshoe crab (1), conifers (1). Those threatened by stone quarrying included: primates (9), millipedes (3), mosses (3), cycads (3), and corals (1). Given that only one species was threatened by both sand extraction and limestone quarrying *(Margaritifera margaritifera*), we re-coded the mining type to sand and gravel extraction for visual representation, as sand extraction is a more direct pressure.

### Global estimate of species impacted by mining

For calculating an approximate total number of animal and plant species impacted by construction mining, we used simple extrapolation methods used by the Intergovernmental Science-Policy Platform on Biodiversity and Ecosystem Services (IPBES)^42, 43^. We calculated the percentage of species threatened with extinction across taxonomic groups and then extrapolated using a mid-low estimate of 8.1 million animal and plant species, of which ca. 5.5 million are insects^20, 79, 80^. Given that the actual number of threatened species is uncertain because it is not known whether Data Deficient (DD) species are actually threatened or not, a range of percentages of species threatened by construction mining was calculated following the IUCN guidelines for reporting on proportion threatened^81^: lower-bound estimate = % threatened assessed species (if all DD species are not threatened); mid-point = % threatened assessed species (if DD species are equally threatened as data sufficient species); upper-bound estimate = % threatened assessed species (if all DD species are threatened). Only global assessments of species (excluding intraspecific taxa) were considered. We followed this approach for three blocks of data.

First, we focused on the Red List comprehensively assessed groups (>80% of species in the group are assessed) containing > 150 species as defined by the Red List reporting approach^38^ (see Figure 2 in https://www.iucnredlist.org/resources/summary-statistics). These criteria led to excluding all insect species. The percentage of species currently threatened with extinction reported as impacted by construction mining averaged 0.27% across groups of animals and plants, excluding insects (**Supplementary Table 2**). Extrapolating by 2.6 million species^80^ there would be 7,060 animal and plant species excluding insects threatened with extinction by construction mining (range = 7,057-7,510 species). However, 74% of species impacted by construction mining fall into poorly-assessed groups and subgroups. For example, the above estimate considers that no gastropods are threatened by construction mining, as the ‘selected group of gastropods’ comprehensively assessed includes exclusively cone snails, of which none were reported as impacted by construction mining. However, construction mining is a well-known threat to snails^36^.

Next, we considered all animals and plants in the Red List together. In this case, the percentage of species currently threatened with extinction and reported as impacted by construction mining was 0.49% excluding insects, and 0.21% for insects. Extrapolating to the number of estimated extant species would result in 24,355 species (range = 19,885-27,754 species) (**Supplementary Table 3**).

Finally, as different taxonomic groups are affected differently by construction mining and are unequally assessed we explored the separate contribution of smaller taxonomic grouping. We considered major groups of animal and plant organisms (following the classification in IUCN Summary Statistics^82^, see Table 1b) containing > 150 species, and averaged for animals and plants excluding insects, and separately for insects. Velvet worms (*n* = 11 assessed species) and horseshoe crabs (*n* = 4) were integrated with ‘Other invertebrates’. Green algae (*n* = 16) and red algae (*n* = 58) were excluded. Insect species were disaggregated by order, with groups of > 150 species, the remaining being combined into ‘Other insects’. On average, 0.50% of the Red List threatened animal and plant species excluding insects and 0.21% insect species are impacted by construction mining. Extrapolating by the number of estimated extant species results in an estimate of 24,246 species [24,234-27,226] (**Supplementary Table 4**).

This estimate should be regarded as a conservative indication of the order of magnitude of the total number of threatened animal and plant species. This approach assumes that undescribed species are neither more nor less likely than described species to be threatened^83^, although they are likely to be more threatened on average as they have narrower distributions than their described relatives^84, 85^. We do, however, recognize that the percentage of threatened species for some of the incompletely evaluated groups might be an overestimate, since assessment efforts might have focused more on species that are likely to be threatened^38^. However, the Red List experts assess either all of a group’s species or choose sets of species to give a representative snapshot of the whole^83^. We also acknowledge that, as with any other threat most species in our database are not uniquely impacted by this mining threat and for some of them other threats might be more influential to their conservation status. Nevertheless, this estimate does not account for the very likely under-reporting of this mining threat within the Red List as suggested by our results. Moreover, Red List data suffers the biases described above and from time-lags in the reporting of threats and extinction events, which makes it slow to identify emerging threats such as construction mining^54, 65, 66^.

## Data availability

Raw data are available from the original sources: https://www.iucnredlist.org and http://datazone.birdlife.org. Economic indicators can be sourced from the World Bank http://databank.worldbank.org The map of global biodiversity hotspots defined by Conservation International is available in https://doi.org/10.5281/zenodo.3261806. Material flow data of non-metallic minerals - construction dominant is available from the Global Material Flows Database of the UN International Resource Panel http://www.resourcepanel.org/global-material-flows-database. The datasets generated and/or analyzed during the current study will be deposited prior publication in the Zenodo Repository.

## Code availability

R scripts used in analysis are available upon request from the corresponding author.

## Acknowledgments

We thank all of the IUCN members and all additional experts who have contributed data and their expertise to Red List assessments or that facilitated the Red List assessments, and those taxonomists and conservationist documenting mining threats to biodiversity. In particular, we thank Olaosebikan Babatunde, Magda Bou Dagher Kharrat, Hederick Dankwa, Hicham Elzein, Craig Hilton-Taylor, Philippe Laleye, Raoul Monsembula, Tilo Nadler, Yenemula Ranga Reddy, Jos Snoeks, and Melanie L Stiassny. The scientific results and conclusions, and any views or opinions expressed herein, are those of the author(s) and do not necessarily reflect those of institutions or data providers. We thank J. Treweek for guidance on good practice for poorly known biodiversity and E.F. Lambin for comments that have improved the work. A.T. received funding from the European Union’s Horizon 2020 research and innovation programme under the Marie Sklodowska-Curie grant agreement No 846474. S.O.S.E.z.E. is supported through NERC’s EnvEast Doctoral Training Partnership (grant NE/L002582/1) in partnership with Balfour Beatty. F.Z.T. was supported by a postdoctoral fellowship from Coordenação de Aperfeiçoamento de Pessoal de Nível Superior (PNPD/CAPES, Finance Code 001). J.L. was supported by the U.S. National Science Foundation and Michigan AgBioResearch.

## Author contributions

A.T. led the research. A.T. and F.F. conceived the study. S.O.S.E.z.E, C.W., F.F., and A.T., screened and coded assessments and performed the literature search. F.Z.T, I.R., F.F., and A.T. conducted the statistical and spatial analyses. A.T. wrote the first draft. All authors discussed the data, analysis and results, and contributed to writing the manuscript.

## Additional information

Supplementary Information is available for this paper. Correspondence and requests for materials should be addressed to A.T.

## Extended Data Figures

**Extended Data Figure 1.**
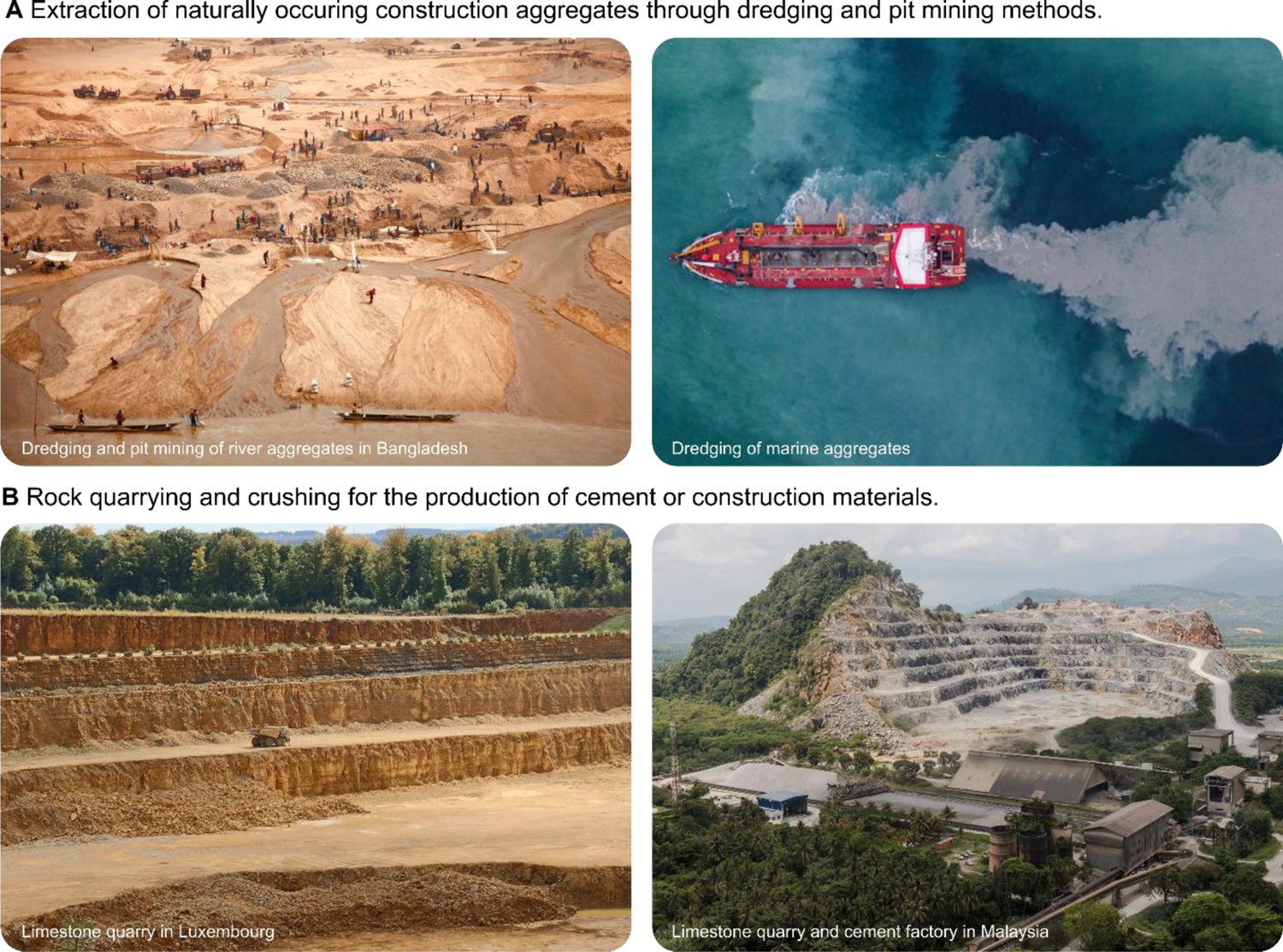
Diversity of construction minerals mining contexts. (**A**) Naturally occurring aggregates are excavated from land deposits (e.g., floodplains, dunes), and dredged from wetlands and shallow coastal waters (top left photo by H. Hayder; top right photo by Sky Cinema). (**B**) Aggregates are also produced by blasting and crushing rocks (e.g., limestone, dolomite, basalt). Limestone is also quarried to produce cement *clinker* or for dimension stones as other rocks (e.g., marble or granite) (bottom left photo by A.J. Johnson; bottom right photo by T. Smith).

**Extended Data Figure 2.**
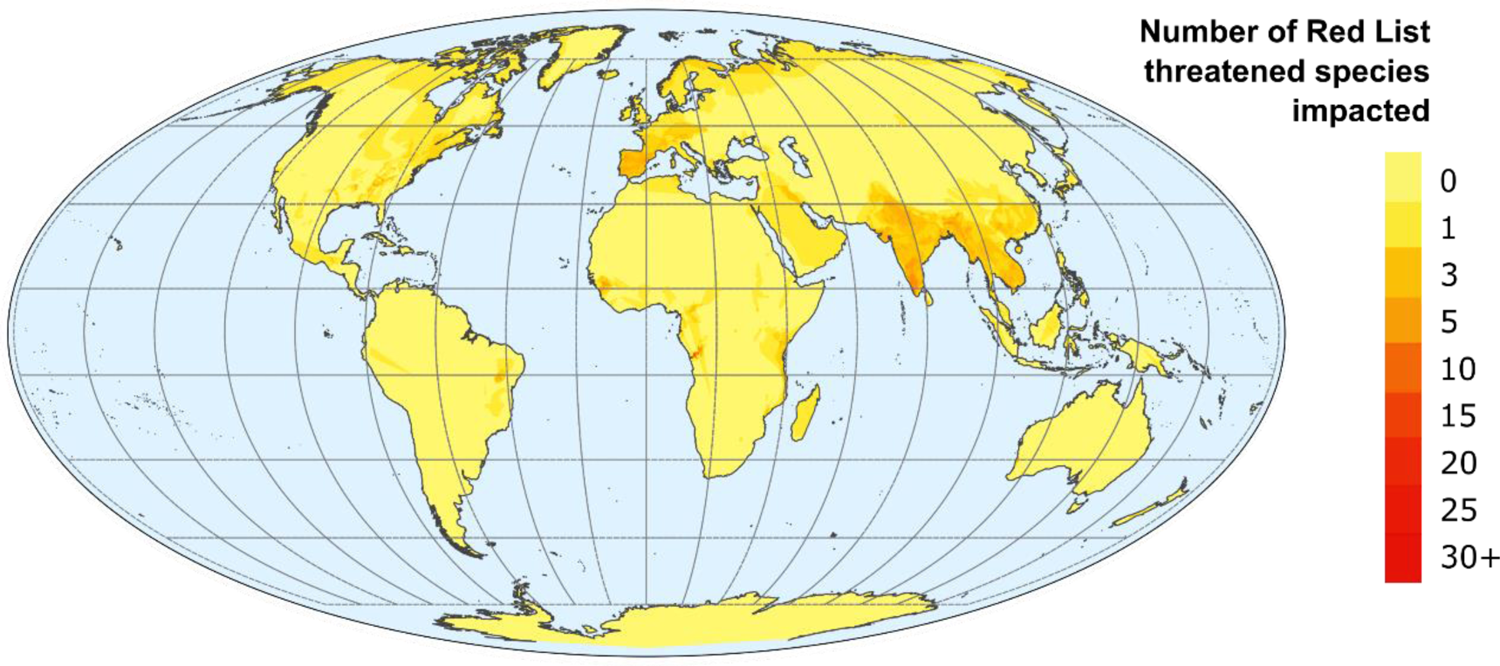
Distribution ranges of the identified Red List threatened species (Critically Endangered, Endangered, or Vulnerable) impacted by construction mining. The actual impact might vary across their range. As virtually no construction mining occurs in the open oceans, the sea surface beyond the continental shelf and at a distance >100 km from the coastline was excluded for clarity.

**Extended Data Figure 3.**
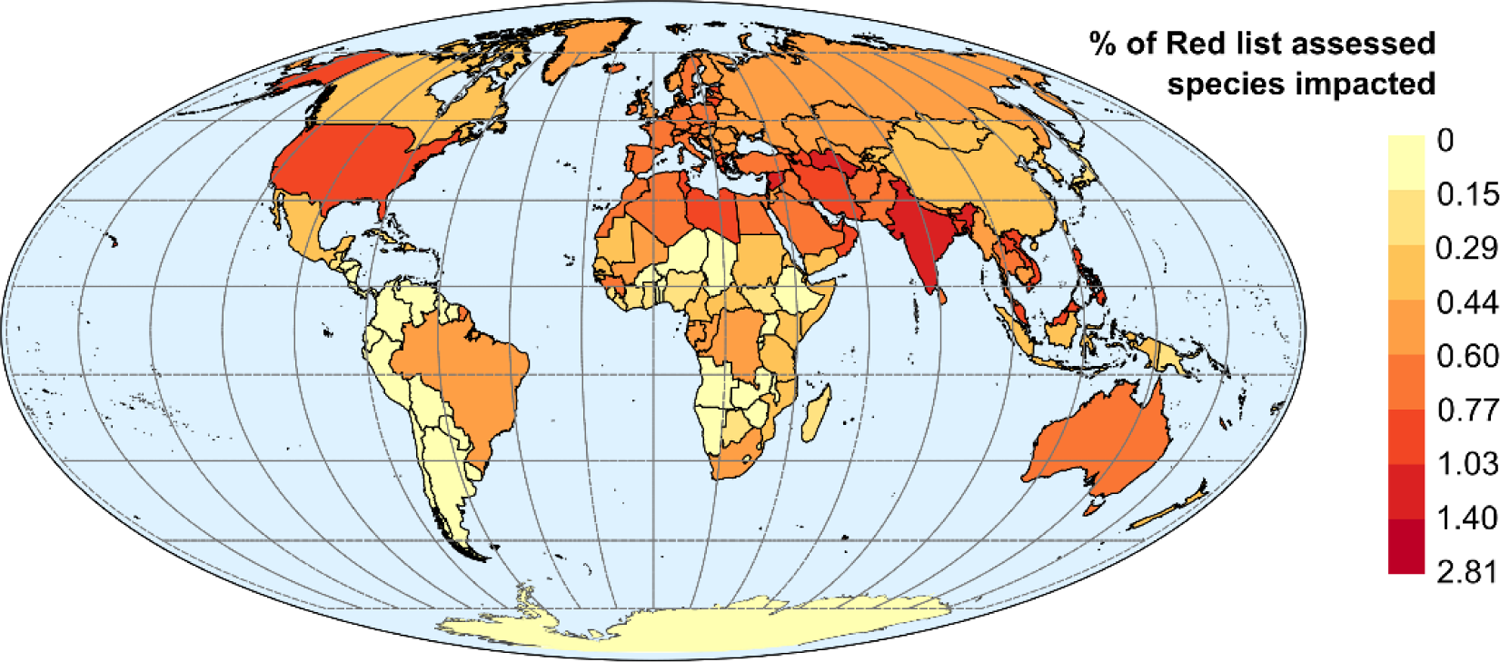
Percentage of Red List assessed (all Red List categories) species impacted by construction mining over the total number of Red List assessed species per country.

**Extended Data Figure 4.**
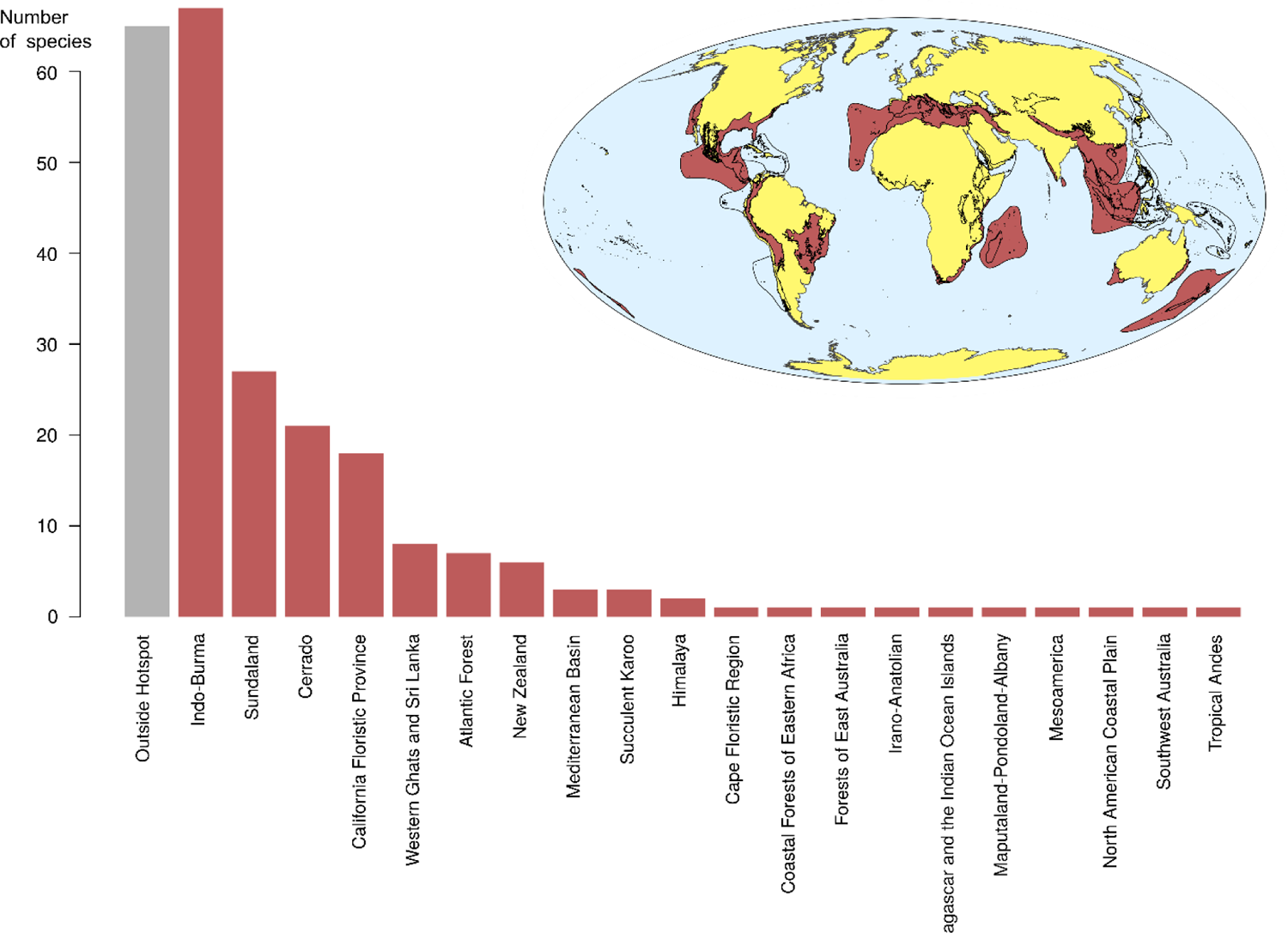
Number of identified new species from species descriptions impacted by construction mining within (red) and outside (gray) biodiversity hotspots. Twenty of the 36 recognized biodiversity hotspots contain at least one observation. We used the freely available Biodiversity Hotspots database version 2016.1^27^ from the Critical Ecosystem Partnership Fund (https://www.cepf.net/our-work/biodiversity-hotspots/hotspots-defined). The map indicates hotspots with (in red) and without (no fill color) observations.

**Extended Data Figure 5.**
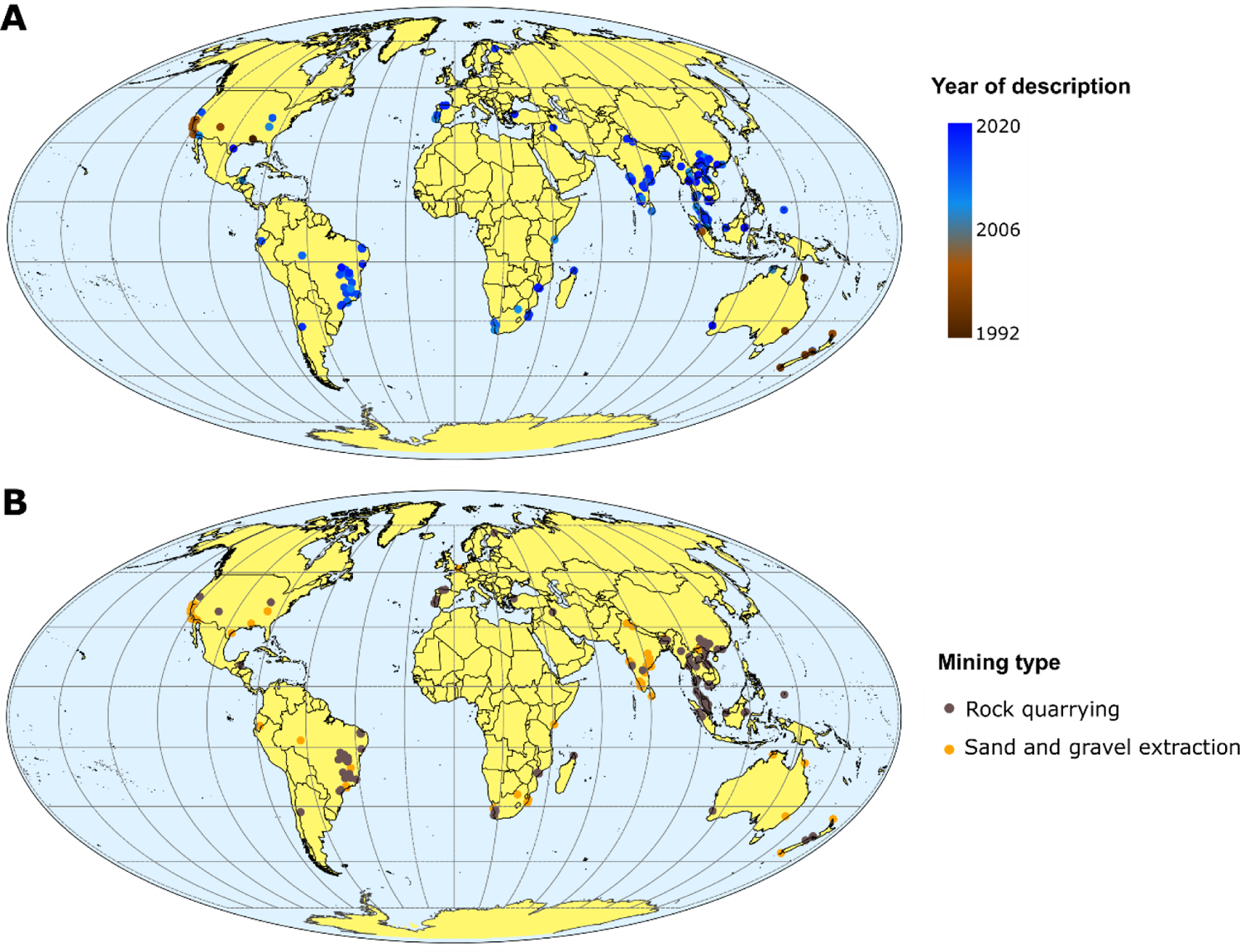
Distribution of identified new species impacted by construction mining by year of description (A), and by type of mining reported as a threat to the species (B).

**Extended Data Figure 6.**
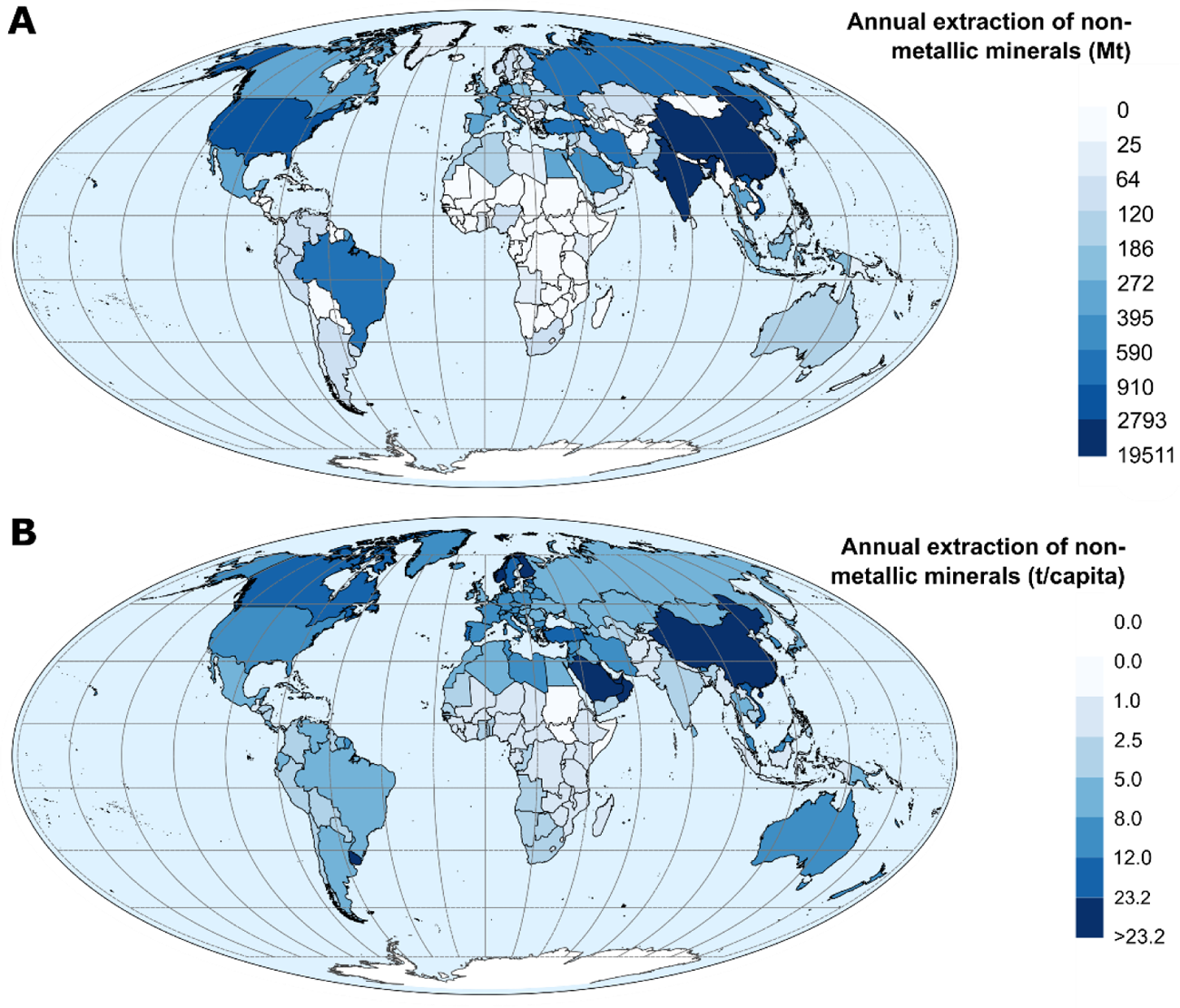
Domestic extraction of non-metallic minerals (mainly construction minerals) from UN and IRP Global Material Flows Database^30^. (**A**) Average annual domestic extraction by country from 2010 to 2017. (**B**) Average annual domestic extraction per capita from 2010 to 2015 (per capita values were not available after 2015).

**Extended Data Figure 7.**
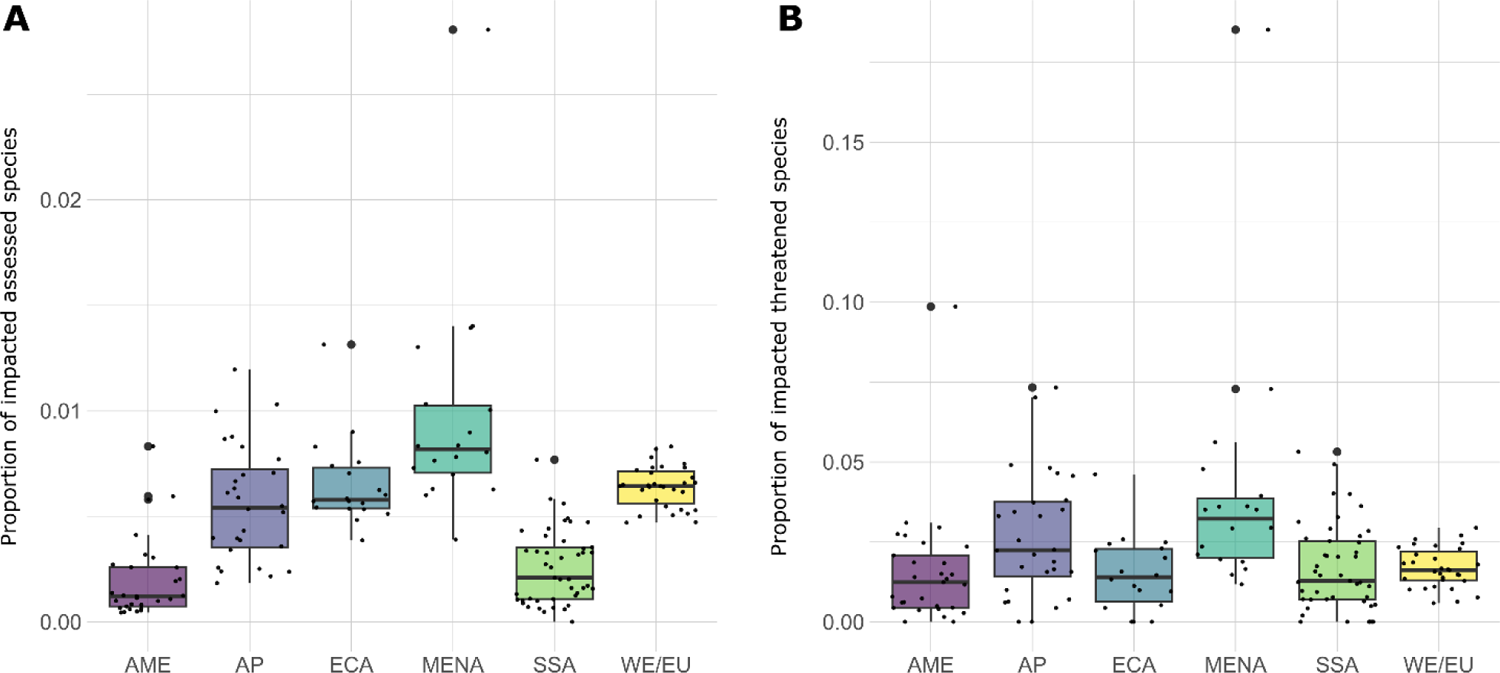
Boxplots of the region and the proportion of species assessed. (A) and threatened with extinction (B) in the Red List that that are impacted by construction mining. Abbreviations: AME: America; AP: Asia-Pacific; ECA: Europe and Central Asia; MENA: Middle East and Northern Africa; SSA: Sub-Saharan Africa; WE/EU: Western Europe / European Union.

**Extended Data Figure 8.**
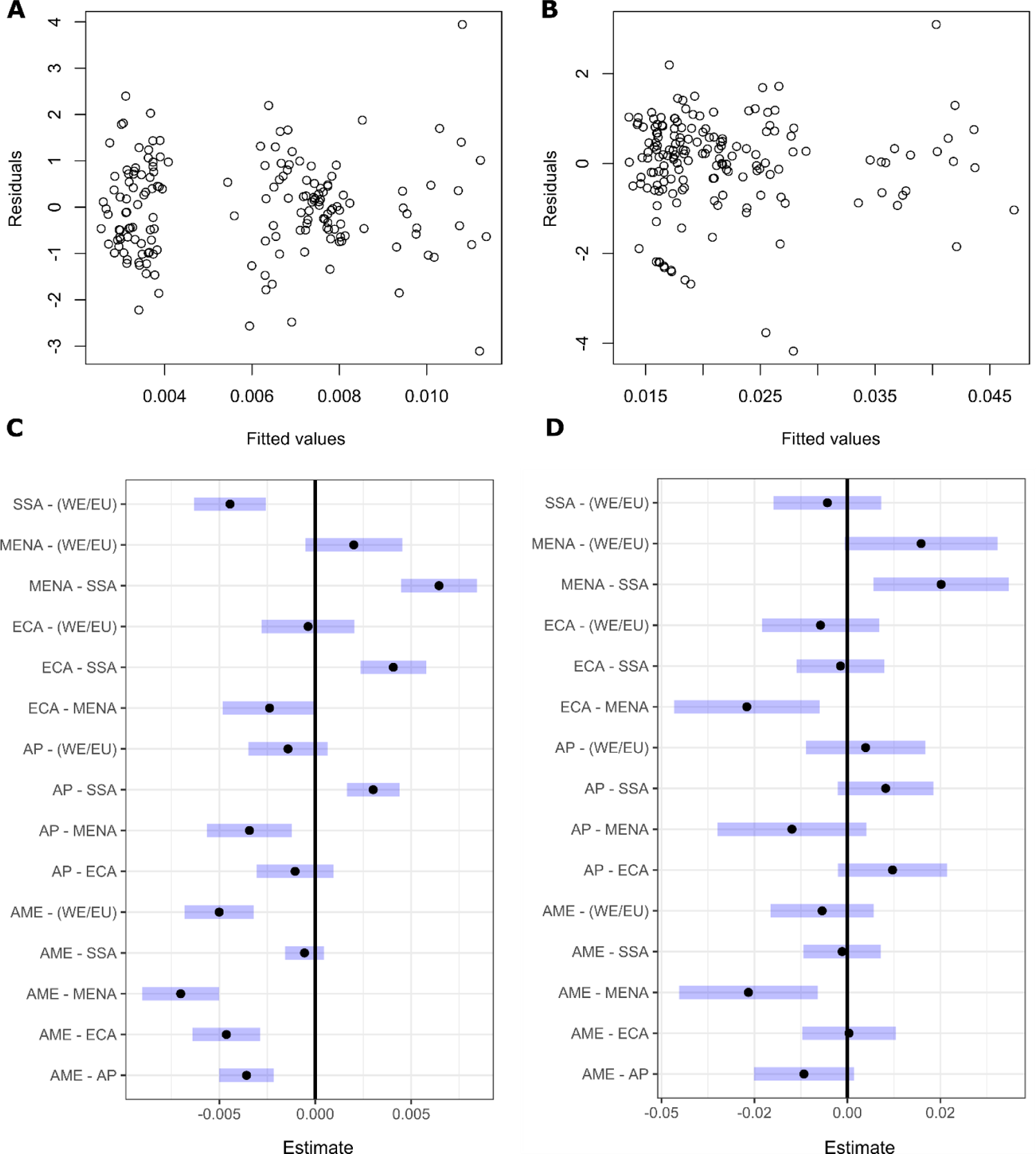
Residual plots of the beta regression models between the proportion of species assessed. (**A**) and threatened with extinction (**B**) in the Red List that are reported as impacted by construction mining and country-level characteristics. Pairwise comparisons among world regions, based on the results of the model with the proportion of species assessed (**C**) and threatened with extinction (**D**). Abbreviations: AME: America; AP: Asia-Pacific; ECA: Europe and Central Asia; MENA: Middle East and Northern Africa; SSA: Sub-Saharan Africa; WE/EU: Western Europe / European Union.

**Extended Data Figure 9.**
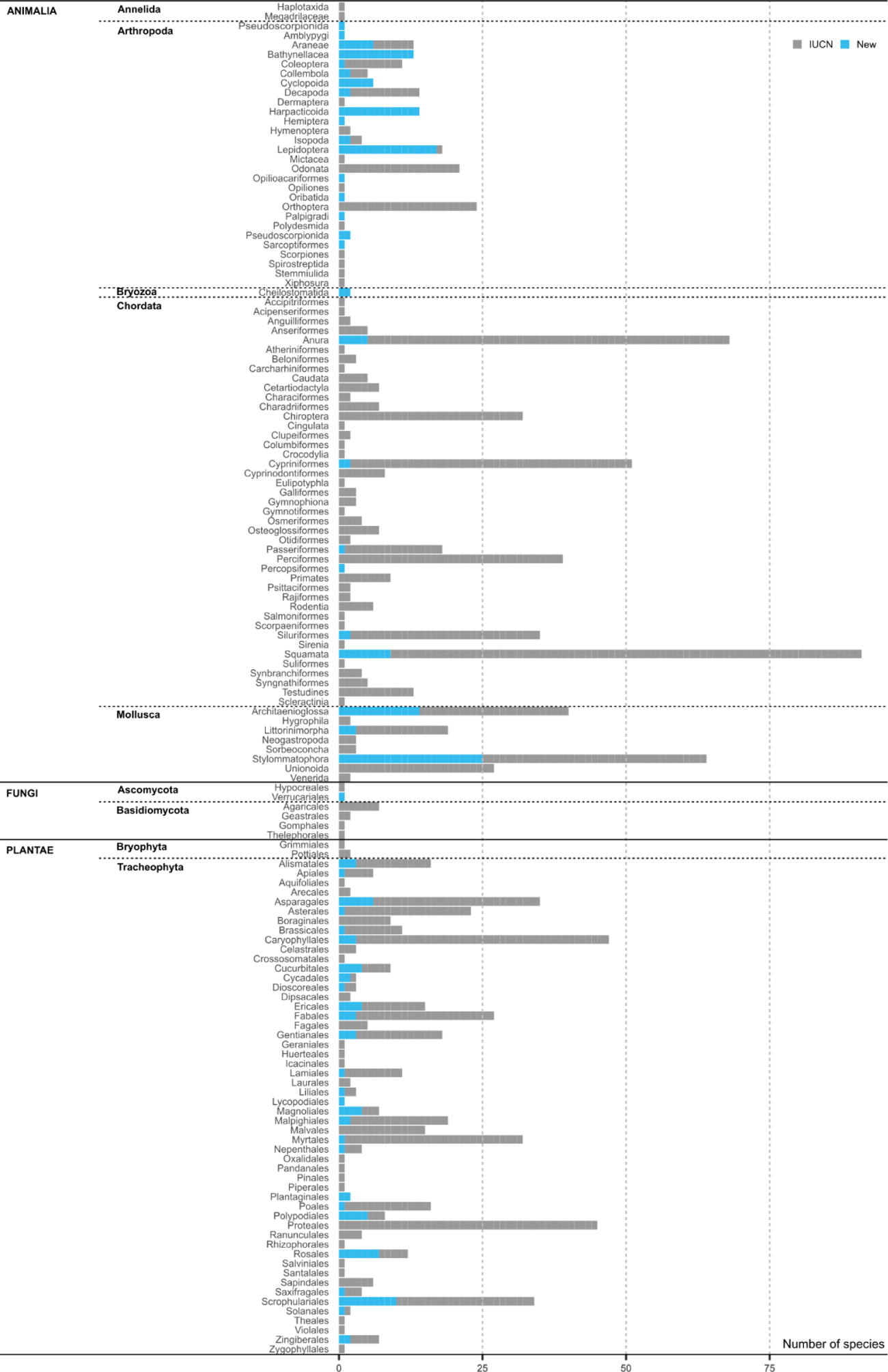
Assessed species and intraspecific taxa of the Red List impacted by construction mining (*n* = 1,066), and new species impacted by construction mining and unassessed by the Red List (*n* = 209), sorted by kingdom and phylum.

## Extended Data Tables

**Extended Data Table 1.** Results of the beta regression model between the proportion of species assessed in the Red List that are reported as impacted by construction minerals mining of the total number of assessed species and country-level characteristics (*P* < 0.001, Pseudo *R*^2^ = 0.60).

| Parameter | Estimate | Standard Error | Z-value | P-value |
| --- | --- | --- | --- | --- |
| (Intercept) | -5.615 | 0.126 | -44.453 | < **0.001** |
| Region - Asia Pacific | 0.796 | 0.111 | 7.197 | < **0.001** |
| Region - Europe and Central Asia | 0.946 | 0.116 | 8.141 | < **0.001** |
| Region - Middle East and Northern Africa | 1.221 | 0.112 | 10.943 | < **0.001** |
| Region - Sub-Saharan Africa | 0.175 | 0.112 | 1.559 | 0.119 |
| Region - Western Europe / European Union | 0.994 | 0.116 | 8.571 | < **0.001** |
| Gross domestic product per capita | $3.6 \cdot 10^{-6}$ | $6.8 \cdot 10^{-4}$ | 1.659 | 0.097 |
| Domestic extraction of non-metallic minerals (t/ha) | $-8.7 \cdot 10^{-5}$ | 0.001 | -0.128 | 0.898 |
| Corruption perception index | -0.006 | 0.003 | -2.300 | **0.021** |

**Extended Data Table 2.** Results of the beta regression model between the proportion of species threatened with extinction in the Red List that are reported as impacted by construction minerals mining of the total number of threatened species and country-level characteristics (*P* < 0.001, Pseudo *R^2^* = 0.14).

| Parameter | Estimate | Standard Error | Z-value | P-value |
| --- | --- | --- | --- | --- |
| (Intercept) | -3.872 | 0.216 | -17.929 | < **0.001** |
| Region - Asia Pacific | 0.462 | 0.184 | 2.508 | **0.012** |
| Region - Europe and Central Asia | -0.022 | 0.222 | -0.100 | 0.920 |
| Region - Middle East and Northern Africa | 0.855 | 0.190 | 4.490 | < **0.001** |
| Region - Sub-Saharan Africa | 0.068 | 0.177 | 0.384 | 0.701 |
| Region - Western Europe / European Union | 0.292 | 0.204 | 1.431 | 0.152 |
| Gross domestic product per capita | $7.7 \cdot 10^{-6}$ | $4.2 \cdot 10^{-6}$ | 1.817 | 0.069 |
| Domestic extraction of non-metallic minerals (t/ha) | -0.001 | 0.001 | -0.477 | 0.633 |
| Corruption perception index | -0.008 | 0.005 | -1.617 | 0.106 |

