## Supplementary Tables 1-4 and Additional Results for "Unearthing the global impact of mining construction minerals on biodiversity"

This document contains:

**Supplementary Tables 1 to 4**

**Supplementary Results**

**Supplementary References**

### Supplementary Tables

**Supplementary Table 1**. Habitat levels of the IUCN habitat classification scheme (version 3.1), habitat classes used in the habitat association analysis (*n* = 33 classes, **Fig. 3**), and habitat classes used for the alluvial diagram (*n* = 16 classes, **Fig. 4**). The number of habitat classes for the alluvial diagram was lower than for the habitat association analysis to ensure a good visualization of the diagram (too cluttered with 33 classes).

| IUCN habitat classification | Habitat classes for alluvial diagram | Habitat class for habitat association analysis | Notes |
|---|---|---|---|
| **1. Forest** | Forest | - | Too broad category for the association analysis |
| 1.1. Forest – Boreal | Forest | Boreal forest | |
| 1.2. Forest – Subarctic | Forest | Boreal forest | |
| 1.3. Forest – Subantarctic | Forest | Boreal forest | |
| 1.4. Forest – Temperate | Forest | Temperate forest | |
| 1.5. Forest – Subtropical/tropical dry | Forest | Tropical and subtropical forest | |
| 1.6. Forest – Subtropical/tropical moist lowland | Forest | Tropical and subtropical forest | |
| 1.7. Forest – Subtropical/tropical mangrove vegetation above high tide level | Forest | Mangroves | |
| 1.8. Forest – Subtropical/tropical swamp | Other wetlands | Freshwater marshes, swamps and peatlands | |
| 1.9. Forest – Subtropical/tropical moist montane | Forest | Tropical and subtropical forest | |
| **2. Savanna** (and subcategories) | Savanna | Savanna | |
| **3. Shrubland** | Shrubland | | Too broad category for the association analysis |
| 3.1. Shrubland – Subarctic | Shrubland | Boreal shrubland | |
| 3.2. Shrubland – Subantarctic | Shrubland | Boreal shrubland | |
| 3.3. Shrubland – Boreal | Shrubland | Boreal shrubland | |
| 3.4. Shrubland –Temperate | Shrubland | Temperate shrubland | |
| 3.5. Shrubland – Subtropical/tropical dry | Shrubland | Tropical and subtropical shrubland | |
| 3.6. Shrubland – Subtropical/tropical moist | Shrubland | Tropical and subtropical shrubland | |
| 3.7. Shrubland – Subtropical/tropical high altitude | Shrubland | Tropical and subtropical shrubland | |
| 3.8. Shrubland – Mediterranean-type shrubby vegetation | Shrubland | Temperate shrubland | |
| **4. Grassland** | Grassland | - | Too broad category for the association analysis |
| 4.1. Grassland – Tundra | Grassland | Tundra | |
| 4.2. Grassland – Subarctic | Grassland | Tundra | |
| 4.3. Grassland – Subantarctic | Grassland | Tundra | |
| 4.4. Grassland – Temperate | Grassland | Temperate grassland | |
| 4.5. Grassland – Subtropical/tropical dry | Grassland | Tropical and subtropical grassland | |
| 4.6. Grassland – Subtropical/tropical seasonally wet/flooded | Grassland | Tropical and subtropical grassland | |
| 4.7. Grassland – Subtropical/tropical high altitude | Grassland | Tropical and subtropical grassland | |
| **5. Wetlands (inland)** | NA | - | Too broad category for the association analysis |
| 5.1. Wetlands (inland) – Permanent rivers/streams/creeks | Rivers and freshwater lakes | Rivers and streams | |

| | | | |
|---|---|---|---|
| 5.2. Wetlands (inland) – Seasonal/intermittent/irregular rivers/streams/creeks | Rivers and freshwater lakes | Rivers and streams | |
| 5.3. Wetlands (inland) – Shrub dominated wetlands | Other wetlands | Freshwater marshes, swamps and peatlands | |
| 5.4. Wetlands (inland) – Bogs, marshes, swamps, fens, peatlands | Other wetlands | Freshwater marshes, swamps and peatlands | |
| 5.5. Wetlands (inland) – Permanent freshwater lakes (over 8 ha) | Rivers and freshwater lakes | Freshwater lakes | |
| 5.6. Wetlands (inland) – Seasonal/intermittent freshwater lakes (over 8 ha) | Rivers and freshwater lakes | Freshwater lakes | |
| 5.7. Wetlands (inland) – Permanent freshwater marshes/pools (under 8 ha) | Other wetlands | Freshwater marshes, swamps and peatlands | |
| 5.8. Wetlands (inland) – Seasonal/intermittent freshwater marshes/pools (under 8 ha) | Other wetlands | Freshwater marshes, swamps and peatlands | |
| 5.9. Wetlands (inland) – Freshwater springs and oases | Rivers and freshwater lakes | Rivers and streams | |
| 5.10. Wetlands (inland) – Tundra wetlands (inc. pools and temporary waters from snowmelt) | Other wetlands | Other wetlands | |
| 5.11. Wetlands (inland) – Alpine wetlands (inc. temporary waters from snowmelt) | Other wetlands | Other wetlands | |
| 5.12. Wetlands (inland) – Geothermal wetlands | Other wetlands | Other wetlands | |
| 5.13. Wetlands (inland) – Permanent inland deltas | Other wetlands | Inland deltas | |
| 5.14. Wetlands (inland) – Permanent saline, brackish or alkaline lakes | Other wetlands | Coastal and saline wetlands | |
| 5.15. Wetlands (inland) – Seasonal/intermittent saline, brackish or alkaline lakes and flats | Other wetlands | Coastal and saline wetlands | |
| 5.16. Wetlands (inland) – Permanent saline, brackish or alkaline marshes/pools | Other wetlands | Coastal and saline wetlands | |
| 5.17. Wetlands (inland) – Seasonal/intermittent saline, brackish or alkaline marshes/pools | Other wetlands | Coastal and saline wetlands | |
| 5.18. Wetlands (inland) – Karst and other subterranean hydrological systems (inland) | Caves | Caves | Caves and other subterranean habitats |
| **6. Rocky Areas** | Rocky areas | Rocky areas | |
| **7. Caves & Subterranean Habitats (non-aquatic)** (and subcategories) | Caves | Caves | Caves and other subterranean habitats |
| **8. Desert** (and subcategories) | Desert | Desert | |
| **9. Marine Neritic** | Neritic habitats | - | Too broad category for the association analysis |
| 9.1. Marine Neritic – Pelagic | Neritic habitats | Deep ocean | |
| 9.2. Marine Neritic – Subtidal rock and rocky reefs | Neritic habitats | Subtidal habitats | |
| 9.3. Marine Neritic – Subtidal loose rock/pebble/gravel | Neritic habitats | Subtidal habitats | |
| 9.4. Marine Neritic – Subtidal sandy | Neritic habitats | Subtidal habitats | |
| 9.5. Marine Neritic – Subtidal sandy-mud | Neritic habitats | Subtidal habitats | |
| 9.6. Marine Neritic – Subtidal muddy | Neritic habitats | Subtidal habitats | |
| 9.7. Marine Neritic – Macroalgal/kelp | Neritic habitats | Macroalgal/kelp | |

| | | | |
|---|---|---|---|
| 9.8. Marine Neritic – Coral Reef (and subcategories) | Neritic habitats | Coral reef | |
| 9.9 Seagrass (Submerged) | Neritic habitats | Seagrass | |
| 9.10 Estuaries | Neritic habitats | Estuaries | |
| **10 Marine Oceanic** (and subcategories) | Deep ocean | Deep ocean | |
| **11 Marine Deep Ocean Floor** (Benthic and Demersal) (and subcategories) | Deep ocean | Deep ocean | |
| **12 Marine Intertidal** | Intertidal zone | - | Too broad category for the association analysis |
| 12.1 Rocky Shoreline | Intertidal zone | Rocky areas | |
| 12.2 Sandy Shoreline and/or Beaches, Sand Bars, Spits, etc. | Intertidal zone | Intertidal shoreline | |
| 12.3 Shingle and/or Pebble Shoreline and/or Beaches | Intertidal zone | Intertidal shoreline | |
| 12.4 Mud Shoreline and Intertidal Mud Flats | Intertidal zone | Intertidal shoreline | |
| 12.5 Salt Marshes (Emergent Grasses) | Intertidal zone | Coastal and saline wetlands | |
| 12.6 Tidepools | Intertidal zone | Intertidal shoreline | |
| 12.7 Mangrove Submerged Roots | Intertidal zone | Intertidal shoreline | |
| **13 Marine Coastal/Supratidal** | NA | - | Too broad category for the association analysis |
| 13.1 Sea Cliffs and Rocky Offshore Islands | Rocky areas | Rocky areas | |
| 13.2 Coastal Caves/Karst | Rocky areas | Rocky areas | |
| 13.3 Coastal Sand Dunes | Coastal dunes | Coastal dunes | |
| 13.4 Coastal Brackish/Saline Lagoons/Marine Lakes | Other wetlands | Coastal and saline wetlands | |
| 13.5 Coastal Freshwater Lakes | Other wetlands | Coastal and saline wetlands | |
| **14 Artificial – Terrestrial** | NA | - | Too broad category for the association analysis |
| 14.1 Arable Land | Farmland and plantations | Arable land | |
| 14.2 Pastureland | Farmland and plantations | Pastureland | |
| 14.3 Plantations | Farmland and plantations | Plantations | |
| 14.4 Rural Gardens | Urban and rural areas | - | |
| 14.5 Urban Areas | Urban and rural areas | Urban areas | |
| 14.6 Subtropical/Tropical Heavily Degraded Former Forest | Forest | Degraded tropical forest | |
| **15 Artificial – Aquatic** (and subcategories) | Human-made wetlands | - | |
| **16 Introduced Vegetation** | - | - | |
| **17 Other** | - | - | |
| **18 Unknown** | - | - | |

[1]http://www.iucnredlist.org/technical-documents/classification-schemes/habitats-classification-scheme-ver3

**Supplementary Table 2**. Estimated number of threatened animal and plant species, excluding insects, impacted by construction mining in 2020. The percentage of threatened species impacted by construction mining was calculated for comprehensively assessed groups (>80% of species evaluated) containing > 150 species as defined by the Red List reporting approach[1]. A range of percentages is provided following the IUCN guidelines for reporting on proportion threatened[2]: lower-bound estimate = % threatened assessed species (if all DD species are not threatened); mid-point = % threatened assessed species (if DD species are equally threatened as data sufficient species); upper-bound estimate = % threatened assessed species (if all DD species are threatened). Red List categories: Extinct in the Wild (EX); Endangered (EN); Critically endangered (CR); Vulnerable (VU); Near threatened (NT); Least concern (LC); and DD.

| | | **Number of species impacted by construction mining** | | | | | | | **Estimated % of threatened species impacted by construction mining[c]** | | |
|---|---|---|---|---|---|---|---|---|---|---|---|
| **Taxonomic group** | **Number of species assessed by 2020** | **EX** | **CR** | **EN** | **VU** | **NT** | **LC** | **DD** | **Lower-bound** (CR+EN+VU)/(total assessed - EX) | **Mid-point** (CR+EN+VU)/(total assessed - EX - DD) | **Upper-bound** (CR+EN+VU+DD)/(total assessed-EX) |
| Amphibians | 7166 | 0 | 2 | 16 | 11 | 11 | 29 | 2 | 0.41 | 0.41 | 0.43 |
| Birds | 11158 | 0 | 5 | 7 | 11 | 8 | 6 | 0 | 0.21 | 0.21 | 0.21 |
| Bony fishes* | 4705 | 0 | 1 | 1 | 0 | 0 | 3 | 1 | 0.04 | 0.04 | 0.06 |
| Cephalopods | 750 | 0 | 0 | 0 | 0 | 0 | 0 | 0 | 0.00 | 0.00 | 0.00 |
| Conifers | 610 | 0 | 0 | 0 | 0 | 0 | 1 | 0 | 0.00 | 0.00 | 0.00 |
| Corals (reef-forming) | 845 | 0 | 0 | 0 | 0 | 0 | 1 | 0 | 0.00 | 0.00 | 0.00 |
| Crustaceans* | 2885 | 0 | 1 | 1 | 0 | 1 | 2 | 0 | 0.07 | 0.07 | 0.07 |
| Cycads | 307 | 0 | 1 | 0 | 0 | 0 | 0 | 0 | 0.33 | 0.33 | 0.33 |
| Dicots* | 4302 | 0 | 12 | 27 | 14 | 12 | 15 | 4 | 1.23 | 1.23 | 1.33 |
| Gastropods* | 633 | 0 | 0 | 0 | 0 | 0 | 1 | 0 | 0.00 | 0.00 | 0.00 |
| Mammals | 5932 | 0 | 1 | 8 | 8 | 12 | 18 | 5 | 0.29 | 0.29 | 0.37 |
| Reptiles* | 342 | 0 | 1 | 1 | 1 | 0 | 0 | 0 | 0.88 | 0.88 | 0.88 |
| Sharks, rays and chimeras | 1186 | 0 | 0 | 1 | 0 | 1 | 0 | 0 | 0.08 | 0.08 | 0.08 |
| *Average* | | | | | | | | | 0.27 | **0.27** | 0.29 |
| *Estimated total number of animal and plant species excluding insects*[a] | | | | | | | | | 7056.92 | **7059.94** | 7509.79 |

*Selected comprehensively assessed subgroups within larger taxonomic groups. Bony fishes include: anchovies; angelfishes; billfishes; blennies; bonefishes; butterflyfishes; cornetfishes; croakers and drums; denticle herring; dragonfishes, lightfishes and relatives; filefishes; ghost pipefishes;

groupers; gulpers, snipe eels and relatives; jacks, pompanos and relatives; ladyfishes; lanternfishes; lizardfishes and allies; pristigasterids; pufferfishes; round herrings; sardines and relatives; seabreams, porgies and picarels; seahorses, pipefishes and relatives; shrimpfishes; sturgeons; Sundaland noodlefishes; surgeonfishes, tangs and unicornfishes; swordfish; tarpons; trumpetfishes; tunas; wolf herrings; and wrasses. Crustaceans include: lobsters; freshwater crabs; freshwater crayfishes; and freshwater shrimps. Dicots include: birches; cacti; magnolias; maples; oaks; protea family; southern beeches; and teas. Gastropods include: cone snails. Reptiles include: marine turtles; seasnakes; chameleons; and crocodiles and alligators; and marine turtles.

[a]Extrapolating by 2.6 million animal and plant species excluding insects[3] as used in the Global assessment report of the Intergovernmental Science-Policy Platform on Biodiversity and Ecosystem Services (IPBES)[4,5]. This estimate of animal and plants species excluding insects follows the assumption that 75% of animal species are insects from Chapman (2009)[6].

**Supplementary Table 3**. Estimated number of threatened animal and plant species impacted by construction mining in 2020. The total percentage of threatened species impacted by construction mining was calculated for animals and plants excluding insects, and exclusively for insects. A range of percentages is provided following the IUCN guidelines for reporting on proportion threatened[2]: lower-bound estimate = % threatened assessed species (if all DD species are not threatened); mid-point = % threatened assessed species (if DD species are equally threatened as data sufficient species); upper-bound estimate = % threatened assessed species (if all DD species are threatened). Red List categories: Extinct in the Wild (EX); Endangered (EN); Critically endangered (CR); Vulnerable (VU); Near threatened (NT); Least concern (LC); and DD.

| Taxonomic group | Number of species assessed by 2020 | Number of species impacted by construction mining: EX | CR | EN | VU | NT | LC | DD | Estimated % of threatened species impacted by construction mining[c]: Lower-bound (CR+EN+VU)/(total assessed - EX) | Mid-point (CR+EN+VU)/(total assessed - EX - DD) | Upper-bound (CR+EN+VU+DD)/(total assessed-EX) |
|---|---|---|---|---|---|---|---|---|---|---|---|
| Animals and plants no insects | 118530 | 3 | 132 | 235 | 212 | 101 | 225 | 40 | 0.40 | 0.49 | 0.52 |
| *Estimated number of species*[a] | | | | | | | | | 10,266.02 | 12,705.19 | 13,578.34 |
| Insects | 10,865 | 1 | 2 | 7 | 14 | 10 | 15 | 5 | 0.18 | 0.21 | 0.26 |
| *Estimated number of species*[b] | | | | | | | | | 9,618.93 | 11,649.32 | 14,175.26 |
| *Estimated total number of animal and plant species*[c] | | | | | | | | | **19,884.94** | **24,354.51** | **27,753.60** |

[a]Extrapolating by 2.6 million animal and plant species excluding insects[3] as used in IPBES Global assessment report[4,5].
[b]Extrapolating by 5.5 million insect species as used in IPBES Global assessment report[4,5], under the assumption that 75% of animal species are insects from Chapman (2009)[6].
[a]Extrapolating by 8.1 million animal and plant species[3] as used in IPBES Global assessment report[4,5].

**Supplementary Table 4**. Estimated number of threatened animal and plant species impacted by construction mining in 2020. The percentage of threatened species impacted by construction mining was calculated for major groups of organisms (see Methods). A range of percentages is provided following the IUCN guidelines for reporting on proportion threatened [2]: lower-bound estimate = % threatened assessed species (if all DD species are not threatened); mid-point = % threatened assessed species (if DD species are equally threatened as data sufficient species); upper-bound estimate = % threatened assessed species (if all DD species are threatened). Red List categories: Extinct in the Wild (EX); Endangered (EN); Critically endangered (CR); Vulnerable (VU); Near threatened (NT); Least concern (LC); and DD.

| | | **Number of species impacted by construction mining** | | | | | | | **Estimated % of threatened species impacted by construction mining[c]** | | |
|---|---|---|---|---|---|---|---|---|---|---|---|
| **Taxonomic group** | **Number of species assessed by 2020** | **EX** | **CR** | **EN** | **VU** | **NT** | **LC** | **DD** | **Lower-bound** (CR+EN+VU)/(total assessed - EX) | **Mid-point** (CR+EN+VU)/(total assessed - EX - DD) | **Upper-bound** (CR+EN+VU+DD)/(total assessed-EX) |
| Amphibians | 7212 | 0 | 2 | 16 | 11 | 11 | 29 | 2 | 0.40 | 0.40 | 0.43 |
| Birds | 11158 | 0 | 5 | 7 | 11 | 8 | 6 | 0 | 0.21 | 0.21 | 0.21 |
| Fishes | 22005 | 0 | 12 | 25 | 40 | 12 | 60 | 9 | 0.35 | 0.35 | 0.39 |
| Mammals | 5940 | 0 | 1 | 8 | 8 | 12 | 18 | 5 | 0.29 | 0.29 | 0.37 |
| Reptiles | 8492 | 0 | 14 | 18 | 21 | 4 | 30 | 7 | 0.62 | 0.63 | 0.71 |
| Arachnids | 358 | 0 | 4 | 4 | 1 | 0 | 0 | 0 | 2.51 | 2.51 | 2.51 |
| Corals | 864 | 0 | 0 | 0 | 0 | 0 | 1 | 0 | 0.00 | 0.00 | 0.00 |
| Crustaceans | 3188 | 0 | 3 | 3 | 5 | 2 | 2 | 0 | 0.35 | 0.35 | 0.35 |
| Mollusks | 8847 | 1 | 28 | 21 | 25 | 12 | 25 | 4 | 0.84 | 0.84 | 0.88 |
| Other invertebrates | 857 | 1 | 1 | 1 | 3 | 2 | 0 | 1 | 0.58 | 0.59 | 0.70 |
| Ferns and relatives | 674 | 0 | 0 | 1 | 0 | 0 | 0 | 0 | 0.15 | 0.15 | 0.15 |
| Flowering plants | 48323 | 1 | 61 | 131 | 87 | 38 | 53 | 12 | 0.58 | 0.58 | 0.60 |
| Gymnosperms | 1016 | 0 | 1 | 0 | 0 | 0 | 1 | 0 | 0.10 | 0.10 | 0.100 |
| Mosses | 282 | 0 | 0 | 0 | 0 | 0 | 0 | 0 | 0.00 | 0.00 | 0.00 |
| *Average* | | | | | | | | | 0.50 | 0.50 | 0.53 |
| *Estimated number of species*[a] | | | | | | | | | 12,948.56 | 12,952.67 | 13,732.48 |
| Coleoptera | 1500 | 0 | 1 | 3 | 2 | 0 | 1 | 3 | 0.40 | 0.40 | 0.60 |
| Hymenoptera | 638 | 0 | 0 | 0 | 0 | 0 | 0 | 0 | 0.00 | 0.00 | 0.00 |
| Lepidoptera | 1254 | 0 | 0 | 1 | 0 | 0 | 0 | 0 | 0.08 | 0.08 | 0.08 |

| | | | | | | | | | | | |
|---|---|---|---|---|---|---|---|---|---|---|---|
| Odonata | 4898 | 0 | 0 | 2 | 5 | 5 | 6 | 2 | 0.14 | 0.14 | 0.18 |
| Orthoptera | 1479 | 0 | 1 | 1 | 7 | 5 | 8 | 0 | 0.61 | 0.61 | 0.61 |
| Other insects | 284 | 1 | 0 | 0 | 0 | 0 | 0 | 0 | 0.00 | 0.00 | 0.00 |
| *Average* | | | | | | | | | 0.21 | 0.21 | 0.25 |
| *Estimated number of species*[b] | | | | | | | | | 11,285.81 | 11,293.70 | 13,493.45 |
| *Estimated total number of animal and plant species*[c] | | | | | | | | | **24,234.37** | **24,246.37** | **27,225.92** |

[a]Extrapolating by 2.6 million animal and plant species excluding insects[3] as used in IPBES Global assessment report[4,5].
[b]Extrapolating by 5.5 million insect species as used in IPBES Global assessment report [4,5], under the assumption that 75% of animal species are insects from Chapman (2009)[6].
[a]Extrapolating by 8.1 million animal and plant species[3] as used in IPBES Global assessment report[4,5]

### Supplementary Results

IUCN Red List species and intraspecific groups impacted by construction mining

*Extinct species*

According to the Red List, by 2020 four species threatened by mining of construction minerals already went extinct. The Saint Helena giant earwig (*Labidura herculeana*) was once the largest earwig in the world, measuring up to 8 cm in length. It became extinct in 2014 following the extraction of stones in its habitat for local construction projects[7,8]. Schmarda's worm (*Tokea orthostichon*) was the first native earthworm formally described in Australasia, found in the Mount Wellington area 150 years ago[9]. This area has been significantly disturbed by urban encroachment and New Zealand’s largest aggregate quarry. A species of peppergrass (*Lepidium obtusatum*), also endemic to New Zealand’s North Island, has not been seen since 1950, with its extinction thought to be partially caused by extraction of its beach-gravel habitat[10]. *Plectostoma sciaphilum* was a land snail found within a single limestone karst that was quarried in 2007 at Bukit Panching in Peninsular Malaysia, destroying the species’ only known habitat[11].

Although the search also identified the cry violet (*Viola cryana*) as extinct due to limestone quarrying and over-collection by botanists[12], such assessment is obsolete. The cry violet is a synonym of the violette de rouen (*Viola hispida*)[13], which is currently listed as critically endangered in the Red List, yet not reported as threatened by mining and quarrying.

*Intraspecific groups*

An additional 19 intraspecific groups (subspecies, varieties, and subpopulations) were identified, of which 11 are threatened with extinction. For example, the Yangtze Finless Porpoise (*Neophocaena asiaeorientalis* subsp. *asiaeorientalis*) and the Ganges River Dolphin (*Platanista gangetica* subsp. *gangetica*) are threatened by river sand mining. The nesting habitat of the North Indian Ocean subpopulation of the green turtle (*Chelonia mydas*) is directly and indirectly threatened by mining and dredging due to habitat destruction and enhanced erosion, and the Dugong’s Nansei subpopulation (*Dugong dugong*) is threatened by coastal mining and land reclamation. A navelwort subspecies (*Omphalodes littoralis* subsp. *gallaecica*) inhabits remaining coastal dune systems in NW Iberian Peninsula (Galicia) and is threatened by coastal sand mining, two rare subspecies of conebush (*Leucadendron elimense* subsp. *elimense* and subsp. *salteri*) inhabiting fynbos in South Africa are threatened by gravel excavation. Finally, a subspecies of saintpaulias from southern Kenya (*Saintpaulia ionantha* subsp. *rupicola*), a subspecies of edible yam endemic to northern Madagascar (*Dioscorea sambiranensis* subsp. *bardotiae*), and the blond-bellied langur (*Trachypithecus pileatus* subsp. *pileatus*) are threatened by limestone mining.

Human uses of Red List assessed species impacted by construction mining

Impacts of mining also cascade into human systems. Many of the Red List assessed species impacted by construction mining provide important functions for ecosystems and people. A quarter of the assessed species and intraspecific groups ($n$ = 268) have identified uses for people according to the IUCN Red List General Use and Trade Classification Scheme (Version 1.0), mostly as human or animal food (13.1%, *n* = 140) or as pets and ornamentals (12.4%, *n* = 132), and of those two thirds (65.3%, *n* = 175) are used for subsistence and livelihood needs.

<u>Conservation actions that are needed for Red List assessed species impacted by construction mining</u>

According to the Conservation Actions Classification Scheme (Version 2.0), the main conservation needs identified for the species and intraspecific groups impacted by construction mining are site and habitat (land and water) protection (58%, *n* = 615 species), site management (36%, *n* = 381), awareness and communication (23%, *n* = 243), ex-situ conservation (e.g., gene-banking, captive breeding; 19%, *n* = 204), and legislation, compliance and enforcement at national and subnational level (15%, *n* = 164).

<u>Research needs of Red List assessed species impacted by construction mining</u>

According to the Research Needed Classification Scheme (Version 2.0), the main research needs identified for the species and intraspecific groups impacted by construction mining are investigating the population size, distribution, and past trends (67%, *n* = 717), followed by life history and ecology (38%, *n* = 408), monitoring of population trends (38%, *n* = 405), and threats (36%, *n* = 381).

13. Kew Science. Viola hispida Lam. *Plants of the World Online* http://powo.science.kew.org/taxon/urn:lsid:ipni.org:names:868315-1.
